# Painting the diversity of a world’s favourite fruit: A next generation catalogue of cultivated bananas

**DOI:** 10.1101/2024.05.29.596104

**Authors:** Julie Sardos, Alberto Cenci, Guillaume Martin, Catherine Breton, Valentin Guignon, Ines Van den Houwe, Yaleidis Mendez, Gabriel L. Sachter-Smith, Rachel Chase, Max Ruas, Ronan Rivallan, Janet Paofa, William Wigmore, David Tilafono Hunter, Angélique D’Hont, Nabila Yahiaoui, Christophe Jenny, Xavier Perrier, Nicolas Roux, Mathieu Rouard

**Author notes:** These authors contributed equally to the study (Julie Sardos, Alberto Cenci).

## Abstract

**Societal impact statement:** Bananas are nutritious fruits of major importance in the tropics and subtropics. Characterizing their diversity is essential to ensure their conservation and use. A catalogue showcasing cultivated bananas genomic diversity was compiled and is to be used as a tool to support the classification of banana cultivars. This research revealed that cultivated banana groups are not all made of identical clones. Materials from recent collecting missions indicated that more banana diversity is expected to be found as the exploration of the banana gene pool continues. These discoveries will drive dynamic conservation strategies for banana genetic resources and will increase their use.

**Summary:**

- Banana is an important food crop cultivated in many tropical and subtropical regions around the world. Due to their low fertility, banana landraces are clonally propagated. However, different factors, such as synonymy and the effects of environment, make their assignment to described sets of clones, or cultivar groups, difficult. Consequently, passport data of accessions in genebanks is often uncomplete and sometimes inaccurate.
- With the recent advances in genomics, a new powerful tool was developed enabling the fine-scale characterization of banana’s ancestry along chromosomes, i.e. *in silico* chromosome painting. We applied this method to a high-throughput genotyping data set obtained from 317 banana accessions spanning most of the known cultivar groups. This set included both genebank and new uncharacterized materials.
- By comparing curated morphological assignation to the genomic patterns resulting from *in silico* chromosome painting, we were able to compile a catalogue referencing the chromosome painting patterns of most of the described cultivar groups.
- Examining the genomic patterns obtained, we discovered intra-cultivar group variability. In some cultivar groups, mitotic recombination or deletions were clonally accumulated in cultivars. In addition, we identified at least 4 cultivar groups in which cultivars likely resulting from distinct sexual events co-existed, notably Pisang Awak in which 5 distinct genomic patterns of two ploidy levels were identified. New patterns were also discovered in the newest materials of the set, showing that a wider diversity of clones still exist *on farm*.

## Introduction

Crop diversity is critical for maintaining the resilience and adaptability of food systems in the face of changing environmental conditions and pests and diseases (Smale and Jamora 2020; McCouch *et al*. 2020). Characterizing this diversity is therefore a much-needed effort to reach a comprehensive overview of the genetic diversity existing within a crop species and to ensure that effective conservation strategies are put in place. This knowledge serves as an essential baseline to monitor the evolution of diversity *in-situ* and to identify new material to be conserved *ex-situ*. Initially, crop characterization consisted in morphological assessments, but it later included molecular descriptions using molecular markers such as RAPD, RFLP, SSR, DArT and SNPs (Powell *et al*. 1996; Agarwal *et al*. 2008; Kilian *et al*. 2012). With the recent progresses made in the fields involving genomics, an unprecedented level of fine-scale characterization can be reached (McCouch *et al*. 2020).

Bananas are an important crop for many tropical and subtropical regions around the world. They are a staple food for millions of people and are a major source of nutrients, income, and employment for many communities. Currently, 80% of global banana production is limited to a few groups of cultivars, such as the dessert Cavendish and the cooking Plantain, but a much wider diversity exists, especially, but not only, in smallholder farms of South Asia and West Oceania, the centre of origin of the crop (Simmonds 1962). This diversity, found in smallholder fields, is conserved through *ex-situ* national, regional and international genebanks, which serve as repositories for landraces, modern varieties and wild relatives and aim at safeguarding as much diversity as possible for present and future generations (Van den houwe *et al*. 2020). Continuous efforts are necessary to properly characterize and rationalize the conserved germplasm and to identify gaps in collections.

The classification of banana cultivars is complex as cultivars are diploid or polyploid hybrids originating from crosses between different wild gene pools. To help in the assignation of cultivars, a scoring method was developed by (Simmonds and Shepherd 1955) on the assumption that most of banana cultivars were derived from the diploid wild species *Musa acuminata* Colla and *M. balbisiana* Colla. By considering the different ploidy levels existing in the crop (diploid, triploid, tetraploid) and the scored relative contribution of both wild ancestors (coded A and B, respectively), Simmonds and Shepherd (1955) introduced the concept of genome constitution groups (e.g. AA, AB, AAA, AAB, ABB). Their work also laid the foundations for the definition of an additional taxonomical level, the subgroups, aiming at refining the classification. These subgroups correspond to cultivars considered to be somatic mutants fixed through vegetative propagation of a single seedling or possibly siblings of related parents (Simmonds 1966; Stover and Simmonds 1987).

After more extensive exploration of banana growing regions and with the beginning of molecular characterization of wild and cultivated germplasm, differential contributions of the subspecies of *M. acuminata* were assessed (Carreel *et al*. 2002; Perrier *et al*. 2011). In addition, other wild ancestors were identified. Notably, it was assessed that some banana cultivars were also hybrids with *M. schizocarpa* (S genome), and with another undefined *Musa* species of the former *Australimusa* section (T genome) (Shepherd and Ferreira 1984; Jarret *et al*. 1992; Carreel *et al*. 1994). Using this classification, a catalogue of the global banana diversity, the Musalogue (Daniells *et al*. 2001), documented 14 genome groups of three ploidy levels, subdivided in 37 subgroups across two botanical sections (Daniells *et al*. 2001; Häkkinen 2013). All subgroups, preferentially called cultivar groups in our study, such as Cavendish, Plantain, Sucrier or Pisang Awak, belong to the *Musa* section.

Despite being visionary for its time, and even with the wide use of published standard descriptors for banana (IPGRI 1996; Taxonomy Advisory Group (TAG) 2016), classifying banana cultivars based on morphology alone is challenging. First, it requires relatively controlled growing conditions since both scores and descriptors were developed in and for *ex-situ* collections and do not consider variations that can be due to environmental conditions (large sense). Second, the identification requires observations at different stages of the plant development, including fructification, requiring space and time as well as suitable climatic growing conditions. Moreover, the assignation of cultivars to cultivar groups depends upon trained eyes and the experience of the observers. Consequently, cultivars are regularly inconsistently classified, with a risk of negative impact on their conservation (Vogel Ely *et al*. 2017). The molecular markers developed later, such as RFLP (Carreel *et al*. 2002), SSR (Hippolyte *et al*. 2012; Christelová *et al*. 2017) or DArT (Risterucci *et al*. 2009; Sardos *et al*. 2016) enabled to characterise the genetic bases of the cultivar groups and helped in the assignation process. However, the use of these technologies for such complex crop do not always allow the unambiguous assignation of accessions.

Recent advances in genomics revealed that cultivar genomes are based on several steps of subgenome combination through hybridization, complicated by homoeologous chromosome exchanges between the genomes of *M. acuminata* and *M. balbisiana* (Baurens *et al*. 2019; Cenci *et al*. 2021; Higgins *et al*. 2023). Moreover, the characterization of the genome diversity of *M. acuminata* subspecies and the fine-scale examination of the A genomes of cultivars showed their mosaic nature. Using a technique known as genome ancestry mosaics painting (or chromosome painting), contributions of *M. acuminata* subspecies could be inferred along the chromosomes of diploids, triploid and tetraploid bananas. This method not only allowed the allocation of specific genomic patterns to cultivars, but it also revealed unidentified ancestors and the underappreciated contribution of *M. schizocarpa* to cultivated bananas (Martin *et al*. 2020; Martin, Cottin, *et al*. 2023).

In this study, we applied the genome ancestry mosaic painting methodology to a large and well curated panel of accessions from the Bioversity International *Musa* Transit Center (ITC) and recent collecting missions. Then, by characterizing the mosaic patterns observed within cultivar groups and for isolated accessions, we developed a catalogue of genomic diversity aiming to be dynamic that will be enriched as banana germplasm characterization expand. This catalogue is to be used as a baseline of reference for further cultivar group assignation for existing and new genebank materials, as well as for *on-farm* projects or any type of work requiring the resolution of banana cultivar classification.

## Materials and methods

### Plant material

A collection of 317 banana accessions was selected for this study, as detailed in **Table S1**. The sources of these materials were diverse: 264 samples were obtained from the Bioversity International *Musa* Transit Centre (ITC) under the form of lyophilized leaves, 44 samples were collected during recent collecting missions, 3 samples were obtained from on-farm projects and 4 reference samples were sampled *in-situ.* Young leaf tissues of samples collected *in-situ* were silica dried on site. A set of 2 samples from the Centre de Ressources Biologiques : Plantes Tropicales (CRB-PT) were obtained as public sequence data from the study of Martin, Cottin, *et al*. (2023).

Significant curation work was undertaken on the taxonomical classification of accessions used in this study to assign or exclude accessions from cultivar groups before genomic characterization. First, passport data available from the *Musa* Germplasm Information System (MGIS-www.crop-diversity.org/mgis) (Ruas *et al*. 2017) was retrieved. Second, and when available, morphological characteristics and taxonomic expert recommendations (Taxonomic Advisory Group members of MusaNet) obtained within the field verification exercise (Chase *et al*. 2014; Van den houwe *et al*. 2020) were checked to correct or refine the classification of some accessions. When no recent field observations were available, collecting mission reports were checked for collector observations in the field. This was notably the case for the collecting mission reports to Papua New Guinea, Cook Islands, Samoa and Tanzania (Arnaud and Horry 1997; De Langhe *et al*. 2001; Byabachwezi *et al*. 2005; Irish *et al*. 2016; Sardos *et al*. 2017; Sachter-Smith *et al*. 2021; Sachter-Smith and Sardos 2021a; b) (**Table S1**).

### DNA Sequencing and genotyping

This study spanned several years and included the preparation and sequencing of accessions in separate batches. As a result, distinct but comparable genotyping technologies were utilized according to the most effective approach at each point in time. For all experiments, DNA from each accession was extracted following a CTAB protocol (modified from Risterucci *et al*. 2000). The libraries for restriction-site-associated DNA sequencing were built with the PstI, or PstI/MseI restriction enzymes, followed by the addition of barcoded adapters, DNA shearing, amplification, and sequencing. The sequencing data were thus generated using either RADseq, ddRADseq or GBS techniques (**Supplementary Table S1**), following the respective protocols established by Davey *et al*. (2010) or Elshire *et al*. (2011). For RADseq, short-insert libraries (300–500 bp) were sequenced to produce 91 bp paired-end reads on an Illumina HiSeq2000 (BGI, Hong Kong, China). For GBS and ddRADseq, libraries were sequenced as 150 bp paired-end reads on an Illumina HiSeq2500 (Genewiz, Azenta Life Sciences, USA) and Illumina NovaSeq 6000 (LGC Genomics GmbH, Germany), respectively.

### Single Nucleotide Polymorphism (SNP) Callings

After demultiplexing with GBSX (Herten *et al*. 2015), FASTQ files (one for each sample) were examined with FastQC. We then used Cutadapt to clean them by eliminating Illumina adapter sequences and trimming low-quality ends with a Phred score > 20 (Martin 2011). Any reads shorter than 30 bp after post-trimming were removed. These reads were subsequently mapped using BWA-MEM (Li and Durbin 2010) to the *Musa acuminata* DH Pahang genome v4 (D’Hont *et al*. 2012; Belser *et al*. 2021), downloaded on the Banana Genome Hub (Droc *et al*. 2022). Re-alignment was done with the IndelRealigner module from GATK v4.1. We then followed the GATK pipeline recommended for a non-model organism by adding the recalibration step. This consisted of performing an initial round of SNP calling on the original uncalibrated data, selecting the SNPs with the highest confidence, and then executing a round of base recalibration on the original mapped reads files. For duplicate samples, a script using Sambamba software was used to merge the recalibrated bam alignment files. The GATK module HaplotypeCaller v4.1 was then used for SNPs and indels calling. Finally, a script gVCF2vcf_gz.pl was written to combine the individual gVCF files obtained into a single VCF file. The GenomicDB procedure from GATK was used to build the gVCF SNP database, containing all the positions, variants and non-variants. The snpcluster exclusion procedure was used to process SNP clusters, set for a threshold of three or more SNPs per 10 bp window. The pipeline used to perform SNP analyses is available at https://github.com/CathyBreton/Genomic_Evolution.

### Genome ancestry mosaic painting

From the resulting VCF files, we used the scripts provided in the VCFHunter version 2.1.2 suite (https://github.com/SouthGreenPlatform/VcfHunter). For each accession, we conserved two alleles by sites with more than 10 reads and less than 1000, minimal frequency >0.05 were discarded (i.e. vcfFilter.1.0.py MinCov:10; MaxCov:1000; minFreq:0.05; MinAl:3; RmAlAlt 1:3:4:5:6:7:8:9:10; RmType SnpCluster). In the next step, we conserved the alleles in common with the alleles identified for 11 ancestral gene pools in Martin, Cottin, *et al*. (2023) (see Identification of ancestry informative alleles), using the vcfSelect.py script. Since this dataset was previously obtained from the whole genome scale, it was possible to intersect genome position of common SNP positions inferred from any genotyping method and mapped on the same reference genome. Then, the allele ratio in individuals was calculated with the allele_ratio_per_acc.py script, generating one file per accessions containing counted allele ratio according to allocated ancestral gene pools (statistics in **Table S1**). When necessary, these files were curated to define the ancestry mosaics of unresolved chromosome segments and to infer potential haplotypes. Finally, genome ancestry mosaics SNP ratios and ancestry allocation were curated and refined, and graphical visualizations were drawn using GeMo (Summo *et al*. 2022). Since the mosaics were inferred using non phased data, the juxtaposition of segments represents introgressions at the position but may not reflect the real haplotype of a given chromosome.

To characterize each cultivar group at molecular level, one accession with genotyping data of good quality and no ambiguous or doubtful classification was selected as a reference (as shown in **Table 1**). Then, patterns of other accessions were compared against the mosaic pattern of this reference accession. During this comparison, certain accessions showed signs of aneuploidy, which could result from *in vitro* conservation processes, especially in the cases of deletions or duplications of chromosomes or chromosome arms (Breton *et al*. 2022). However, these aspects of aneuploidy are not elaborated upon in this study or represented graphically in the catalogue (**Dataset S1**). Other events, usually smaller in size and repeated in several accessions, such as small duplications and deletions, were considered ancestral events that accounted for the creation of different patterns, as described in Martin, Cottin, *et al*. (2023).

**Table 1.**
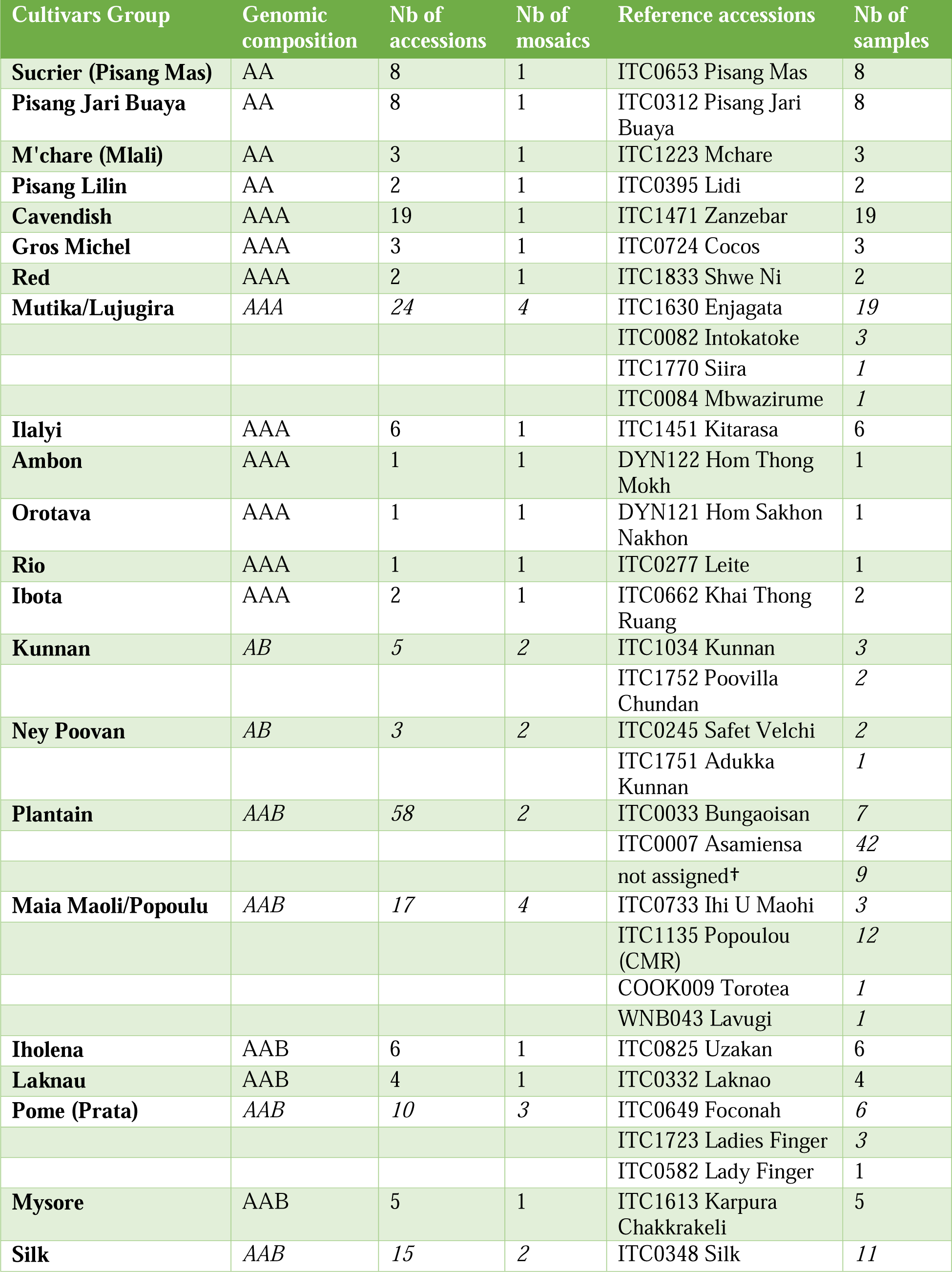
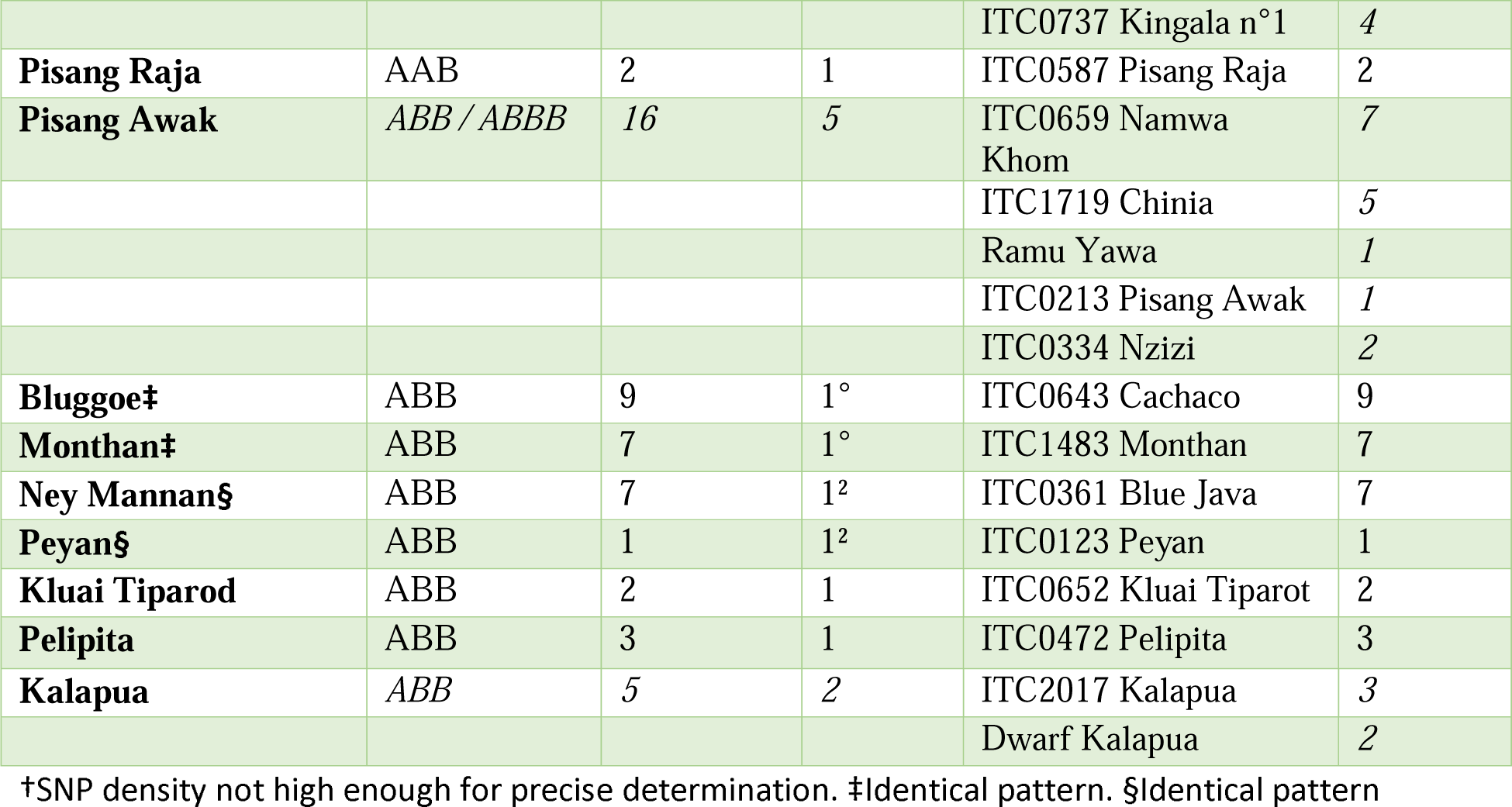
Overview of banana cultivar groups and their genomic compositions, alongside the number of accessions and mosaic patterns identified for each cultivar group. It also includes reference accession names with their identifiers and the total number of samples assigned to each cultivar group.

## Results

In our study, we carefully analysed 317 accessions, through genome ancestry mosaic painting, revealing genomic mosaic patterns at the chromosome level for each cultivated banana. Comparisons of patterns, combined with the curated taxonomical assignation of each accession, enabled the identification of reference patterns for cultivar groups as morphologically defined in the Musalogue (Daniells *et al*. 2001) and in De Langhe *et al*. (2001) for the Ilalyi group. Additionally, our analysis identified accessions that did not match with any cultivar group, both morphologically and genomically. If these unclassified accessions had unique genomic patterns or matched only with accessions known to be synonyms, i.e. same cultivars with different names, we treated them as individual accessions. If the same genomic pattern appeared in several accessions that were not synonyms, we considered them as clusters of morphological variants. The cultivar group patterns, clusters of accessions, and specific genomic patterns of individual accessions were organized and compiled into a catalogue (**Dataset S1**), as exemplified in **Fig. 1**. Nine remaining patterns for which morphological characterization is still ongoing were presented separately (**Dataset S2**).

**Fig. 1.**
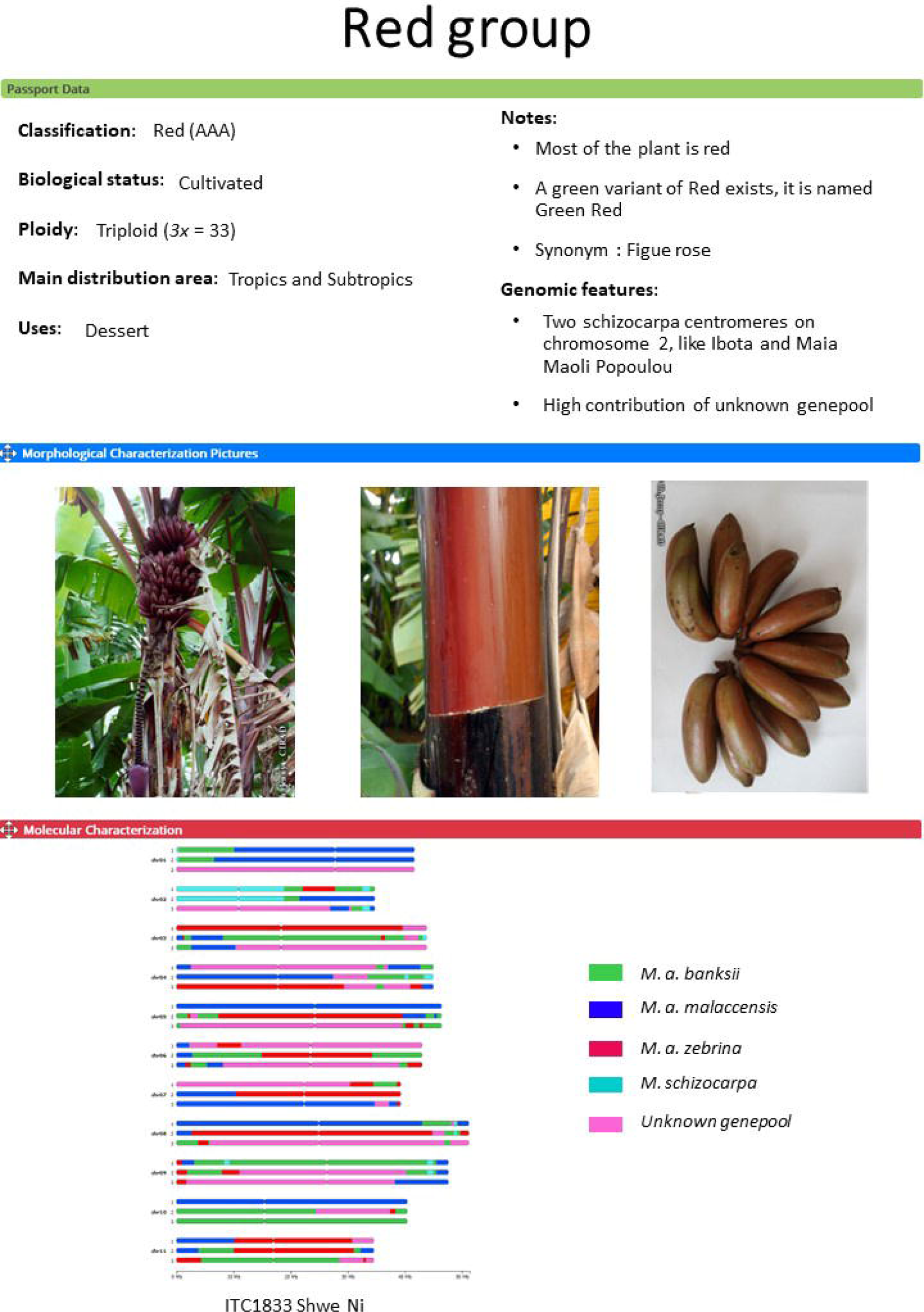
Example of the Red cultivar group in the catalogue. Each entry is divided into 3 sections: Passport data, Pictures and Molecular characterization. The Passport data section includes basic information as the biological status, ploidy, main distribution area and uses with notes on the cultivar group and genomic features. Up to 3 pictures are intended to be representative of the cultivar group. The molecular characterization contains from 1 to 5 mosaic patterns with the name of the reference accession(s) used for the mosaics painting. Coloured segments show the contribution of each of the ancestral genepools (Martin, Cottin, et al. 2023).

Overall, the cultivar groups were well differentiated, and their genetic backgrounds were sufficiently discriminating to assign accessions to specific cultivar groups (**Table 1**). We identified a total of 83 unique mosaic patterns. These patterns corresponded to 31 previously defined cultivar groups (46 patterns; 256 accessions) and to 61 additional accessions or clusters of accessions (37 patterns). Out of the 27 cultivar groups for which more than one accession was available, 18 were homogeneous, i.e. they were composed of accessions with strictly identical genomic mosaics. Conversely, the 9 other cultivar groups displayed heterogeneity, i.e. they were composed of accessions with several mosaic patterns.

We observed that the genomic mosaics of the described cultivar groups exhibited a range of ancestral contributions (**Table 2**). The *M. acuminata* ssp. *banksii,* extended with the accessions ‘Agutay’ (ssp. *errans)* and ‘borneo’ (ssp. *microcarpa)* (referred to as banksii) is the only ancestral genepool for which centromeres were always present, with a minimum of 2 centromeres being observed in 7 patterns. *Musa acuminata* ssp. *zebrina* (referred to as zebrina) and *M. schizocarpa* (referred to as schizocarpa) were consistently present across cultivar groups at least under the form of introgressions, corroborating the findings reported by Martin, Cottin, *et al*. (2023). However notable exceptions were observed in the cultivar group Klue Teparod (ABB) from mainland South-East Asia and the ‘Auko’ clones (ABB) from Papua New Guinea. *M. acuminata* ssp. *malaccensis* (referred to as malaccensis) was also present in 35 patterns (24 cultivar groups). Then, the presence of previously unknown genepools, referred to as m1 and proposed to be *M. acuminata* ssp. *halabanensis* (referred to as halabanensis) in Martin, Cottin, *et al*. (2023), and m2 (referred to as unknown) were present in 14 (11 cultivar groups) and 18 patterns (15 cultivar groups) respectively. *Musa acuminata* ssp. *burmannica,* including ssp. s*iamea* (referred to as burmannica) was found to contribute only to the Klue Teparod cultivar group, along with banksii and schizocarpa for the A haplotype. The B genome contributor, *M. balbisiana,* (referred to as balbisiana) was included in 2/3 of the patterns within cultivar groups, with a minimum of 10 centromeres in 3 triploid cultivar groups and a maximum of 34 centromeres in the pattern Pisang Awak-4x-3. These proportions were in general in line with expectations based on the genomic constitution AB, AAB, or ABB. However, and as noted in Cenci *et al*. (2021), its proportion varied from strict 1:2, 1:3, 2:3 or 3:4, considering the presence of homoeologous exchanges between the A and B genomes. Several of the individual accessions exhibited very distinctive patterns. It included accessions with a notable contribution from burmannica, or cultivars with high contribution of zebrina, as well as tetraploids that exhibited a complete haplotype of Australimusa (T) genome, now included in the former Callimusa section (Häkkinen 2013).

**Table 2.**
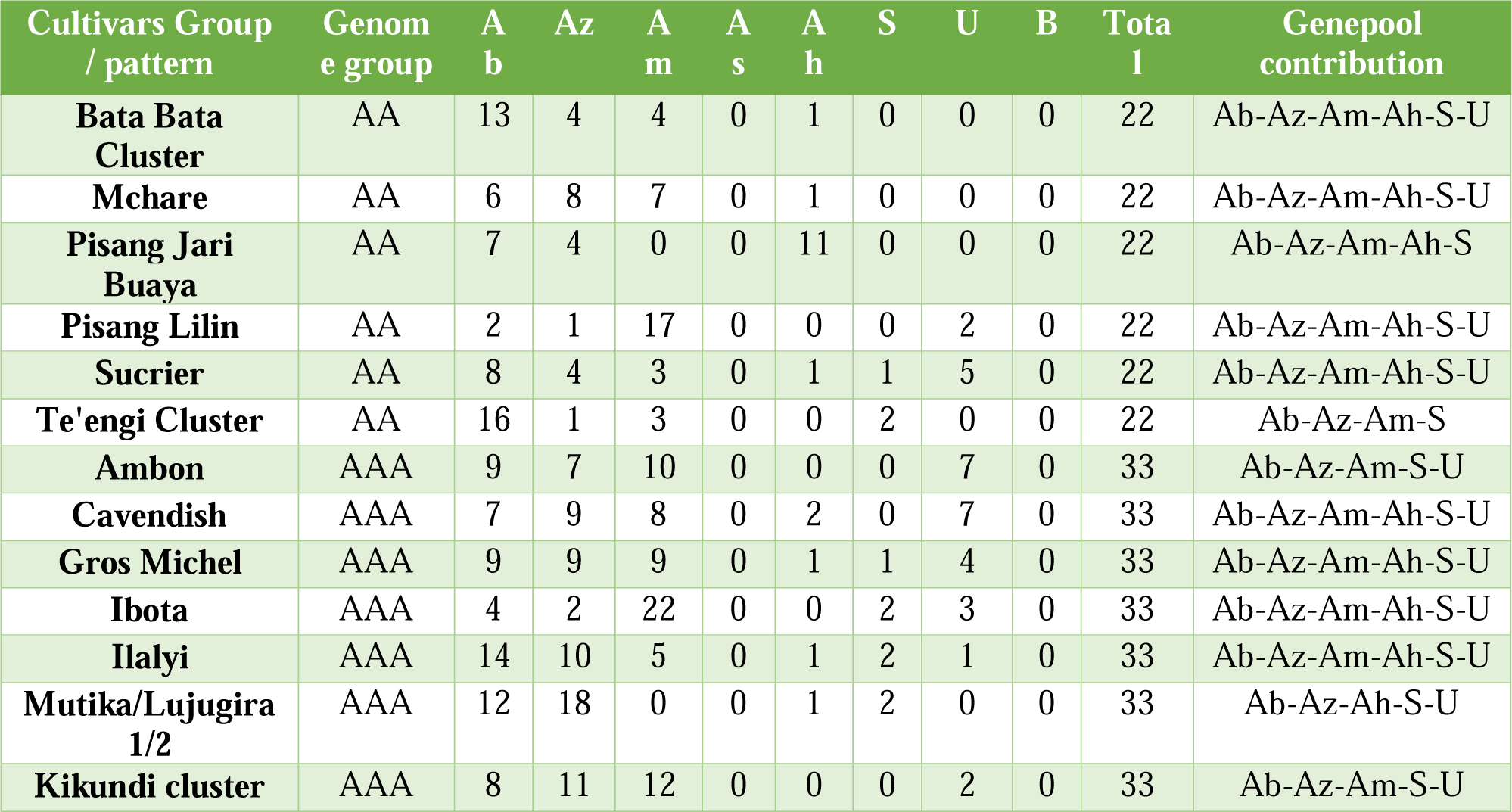
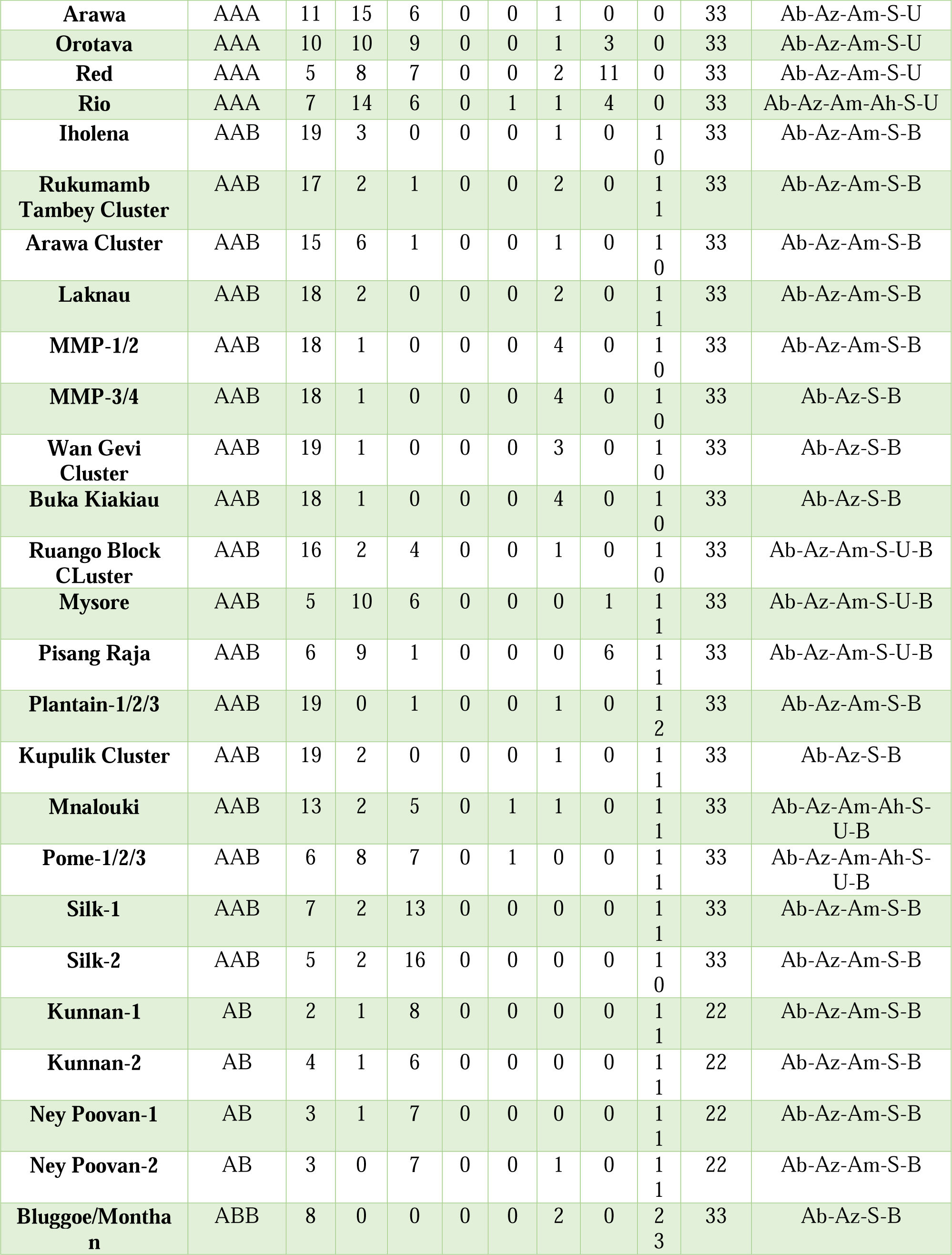
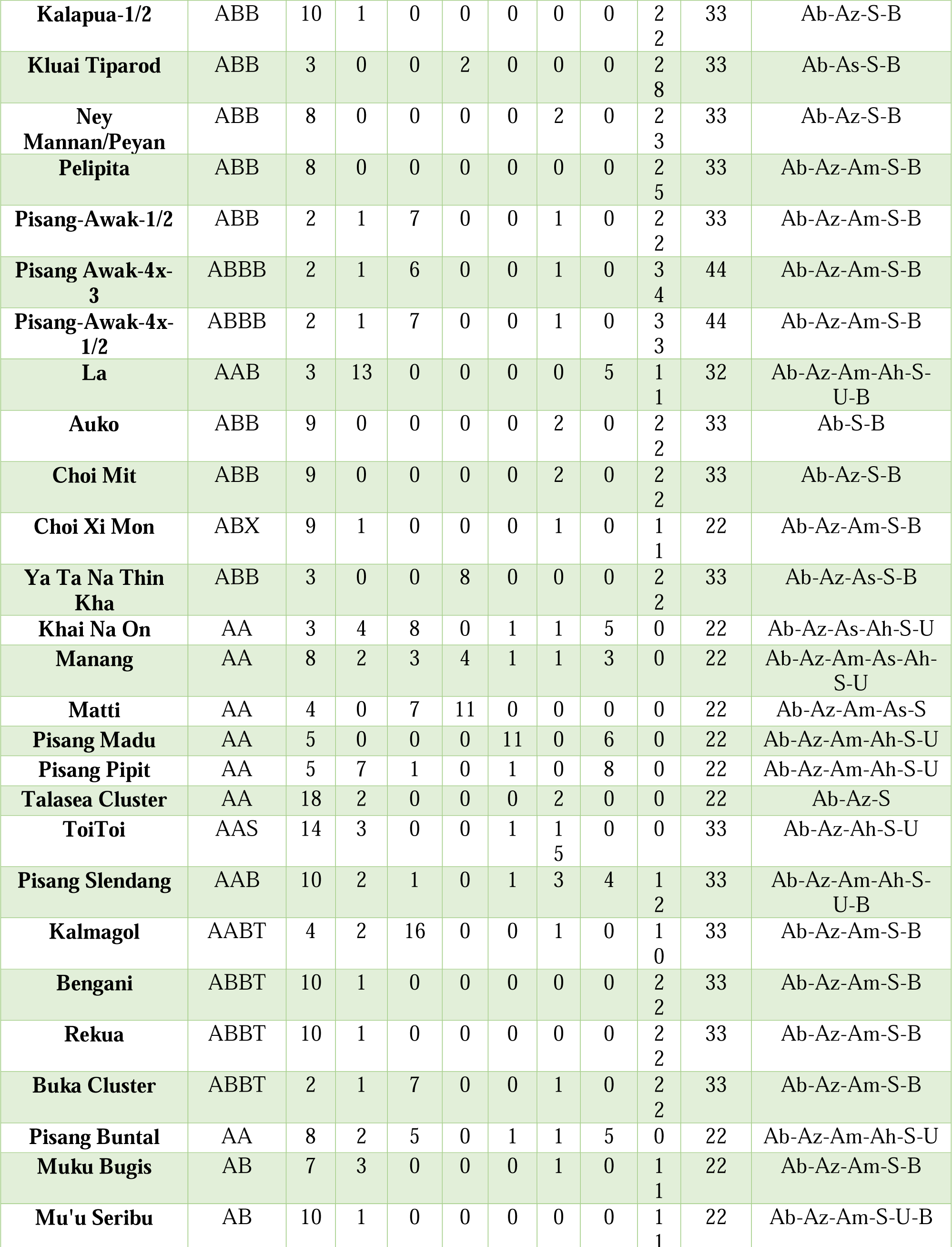
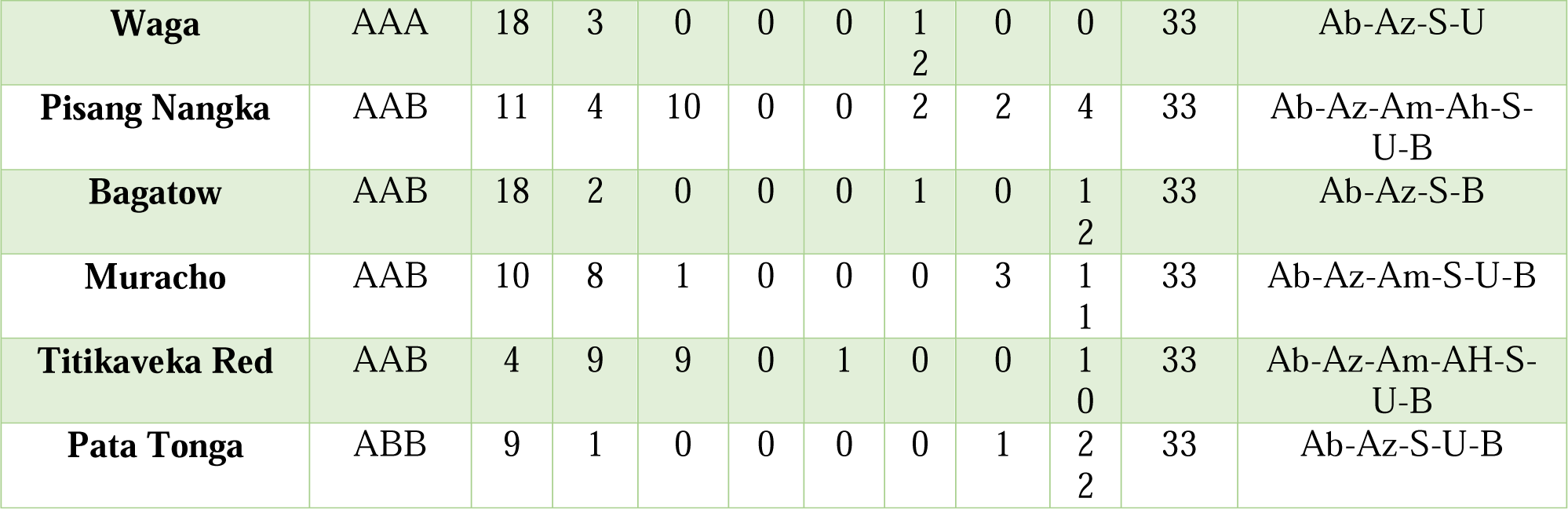
Genepool contributions to various banana cultivar groups and mosaic patterns, detailing the number of centromeres contributed by each genetic source based on the reference chromosome structure. The contributors are denoted as follows: Ab for *banksii*, Az for *zebrina*, Am for *malaccensis*, As for *burmannica/siamea*, Ah for*. halabanensis*, S for *schizocarpa*, U for unknown, and B for *balbisiana*.

### Homogeneous cultivar groups

Homogeneity was observed across cultivar groups of various ploidy levels and genomic compositions.

#### Diploid cultivar groups

With an AA genomic constitution, Pisang Jari Buaya (comprising 7 accessions) and Sucrier (8 accessions) were homogenous in their mosaic patterns. The patterns of the two accessions classified as Mchare were identical, as well as for the two accessions identified as Pisang Lilin.

#### Triploid cultivar groups

In the triploid cultivar groups with an AAA genomic composition, no variation was identified within the 18 accessions of Cavendish analysed. Similarly,, Gros Michel (3 accessions), Red (2 accessions) and Ibota (2 accessions) were homogeneous. After curation, we identified 6 accessions from Tanzania wrongly assigned to the Mutika/Lujugira group that corresponded to the Ilalyi group as described by De Langhe et al. 2001 and genetically validated in Perrier *et al*. (2019). We therefore revived the Ilalyi group that was previously removed from the passport data. These accessions shared the same mosaic pattern, with an ancestral basis composed of a combination of zebrina and banksii similar to the one observed in the Mutika/Lujugira group, but with an additional important contribution of malaccensis (5 centromeres) (**Table 2**). Lastly, due to the availability of only one accession for each of the cultivar groups Ambon, Rio, and Orotava, it was not possible to investigate potential variations within their respective mosaic patterns. The study would benefit from more samples to confirm their monoclonal status.

In the triploid cultivar groups with an AAB genomic composition, Laknau (4 accessions), Mysore (5 accessions), Pisang Raja (2 accessions) and Iholena (6 accessions) were homogenous.

In the triploid cultivar groups with an ABB genomic composition, the Pelipita (3 accessions) and Klue Teparod (2 accessions) groups were genetically uniform while a wider sampling remains necessary to validate this observation. Finally, we identified 2 patterns corresponding to 4 cultivar groups with a shared A genome background. The Bluggoe group (10 accessions) and the Monthan group (5 accessions) exhibited identical genetic mosaic patterns, despite differences in morphologies. For instance, Bluggoe fruits are mostly straight and horizontal or slightly erect, while Monthan’s curve upwards. Similarly, Ney Mannan (7 accessions) and Peyan (1 accession) groups also shared the same mosaic pattern, although we noted slight differences in the levels of heterozygosity within the *M. balbisiana* haplotypes. A larger sample of Peyan representatives and better discrimination of allelic diversity in the B genome would be necessary to provide clearer insights.

### Heterogeneous cultivar groups

Two types of heterogeneous cultivar groups could be distinguished. The first type, which includes cultivar groups such as Plantain and Mutika/Lujugira, displayed mosaic patterns differentiated by only small chromosomal region showing variations in allelic ratio. In the second type of heterogeneous cultivar groups, we found two or more mosaic patterns, each exhibiting multiple differences likely resulting from different mechanisms of diversification.

#### Mutika/Lujugira

Mutika/Lujugira is a triploid cultivar group with a AAA genomic constitution that is typical to Burundi, Uganda, Democratic Republic of Congo, Cameroon, Kenya, Rwanda and Tanzania. For this cultivar group, we conducted an important curation of the passport data, notably by consulting the collecting mission reports when available. In this set of 34 AAA accessions from East Africa, we identified four nearly identical mosaics corresponding to 24 accessions, including well described Mutika/Lujugira cultivars such as the popular ‘Mbwazirume’ (Shepherd 1957). These mosaics’ main contributors were zebrina, banksii and schizocarpa. A pattern variation was observed in ‘Guineo’, ‘Intokatoke’ and ‘Makara’ in which a small interstitial region of the second arm of chromosome 10 displayed a mitotic homologous exchange between one of the zebrina haplotypes and the banksii haplotype. In addition, and on the same chromosomal region, the ‘Siira’ accession exhibited a small deletion on the banksii haplotype. Finally, the accession ‘Mbwazirume’ also appeared to be a variant with a switch of the allelic ration resulting from a mitotic recombination on the first telomere of chromsome 10 (**Fig. S1**). No correlation was found between these variations and the proposed clonesets from Karamura *et al*. (2010). Finally, three additional mosaics discovered in this set were not assigned to Mutika/Lujugira but corresponded to Tanzanian accessions from the homogeneous Ilalyi group described earlier.

#### Plantain

The 58 Plantain accessions of the sample were remarkably homogeneous, but a small variation was detected on chromosome 10 between ∼8.5 Mb and 13 Mb. We interpreted this change as a diploid region (balbisiana – banksii) resulting from a small deletion of one of the banksii haplotypes present in the original pattern. This predominant variation was detected in 42 accessions out of 49, for which this region could be characterized. Interestingly, the 4 accessions with origins in Asia, ‘Bungaoisan’ (a medium French) and ‘Daluyao’ (Medium True Horn) from the Philippines, as well as ‘Mantreken’ from Indonesia and ‘Nendran’ (French) from India all had the original mosaic pattern without deletion. The three other accessions with a mosaic without the deletion were ‘Agbagba” from Nigeria (a medium false-horn Plantain widely cultivated in West Africa according to Adheka *et al*. 2013), ‘Big Ebanga’ (a giant false horn from Cameroon, possibly synonym of Agbagba), and Maiden Plantain (a French Plantain received from Honduras but of unknown origin)

#### Pome

Pome cultivars are AAB triploids that originated in India and are now very popular in Brazil and Hawaii under the name Prata. Ten accessions of the sample displayed a mosaic pattern associated with the Pome group, within which three pattern variations were observed. The first one, identified in 6 accessions from different countries, shows one deviation from a pure AAB pattern, i.e. an A/B recombination (A3:B0) on the first telomere of chromosome 3. The second one, present in 3 Pome accessions received recently from India, also shows an A/B recombination (A3:B0) but in the interstitial region of the first arm of chromosome 9. This recombination is also present in the third pattern, in addition to another one on the first telomere of chromosome 10 **(Dataset S1)**. This last pattern was identified in one accession from Australia, ‘Lady Finger (Nelson)’, sometimes referred to as belonging to the Nadan group in other collections and which is tetraploid for chromosomes 8. Except for the extra chromosome 8 of Pome 3 which has an *M. acuminata* – *M. schizocarpa* ancestry, the variations observed between the three patterns are linked to A-donor introgressions in B chromosomes. Since all these introgressions correspond to genepools also present in one of the two A genomes, it is difficult to assess whether the three Pome patterns were derived clonally from each other or were obtained through different sexual events. However, the banksii introgression observed on the B chromosome 9 of Pome-2 and Pome-3 occurs frequently in cultivars of AB, AAB or ABB genomic constitutions, suggesting a common ancestry. This pattern is more likely to have been inherited sexually rather than arising from a new, independent mitotic recombination. The observed variations may have resulted from a combination of clonal diversification and sexual events (at least two from similar parents).

#### Maia Maoli/Popoulu (MMP)

After curation of passport data and a morphological trait check, we identified 16 accessions affiliated to the Maia Maoli/Popoulu group corresponding to 4 genomic patterns. The Maia Maoli/Popoulou patterns, with an AAB genomic composition, are characterized by contributions from banksii, schizocarpa, and zebrina for the A genome, with little to no presence of malaccensis. A striking feature of these four patterns is the absence of B centromere and the presence of two S centromeres on chromosome 2 (**Table 2**, **Dataset S1**). Out of the 16 accessions, MMP-2 was the most frequent pattern with 12 representatives from both Polynesia (Cook Islands, Samoa and Hawaii) and Melanesia (Bougainville and New Britain Islands in Papua New Guinea). Three accessions from Cook Islands, Tahiti and Samoa exhibited the pattern MMP-1. The differences observed between MMP-1 and MMP-2 are slight. The pattern MMP-1 has a small balbisiana introgression in the interstitial region of the first arm of chromosome 1, and a duplication of the balbisiana first telomere on chromosome 7. The patterns MMP-3 and MMP-4 were identified in one accession each, ‘Mango Torotea’ from Cook Islands and ‘Lavugi’ from New Britain Island, respectively. These patterns are significantly different from MMP-1 and MMP-2 with more than 10 discriminating regions each. They also differ from each other by 15 events. If MMP-1 and MMP-2 may be derived from each other by clonal diversification, this is likely not the case of MMP-3 and MMP-4, which may have resulted from independent sexual events with similar parental contribution.

#### Silk

Two closely related patterns were detected in the 15 accessions classified as Silk confirmed previous findings (Sardos *et al*. 2016). Eleven accessions were displaying the pattern Silk-1 while 4 displayed the pattern Silk-2. Interestingly, the Silk-2 accessions were all collected in Africa (Burundi, Tanzania, Congo). The differences observed between the two Silk genomic mosaic patterns were important. Notably, 5 of their 33 centromeres were of different origins. For example, on chromosome 5 both Silk groups display one B chromosome, but Silk-1 displays two banksii centromeres while Silk-2 displays one banksii and one zebrina centromere (**Table 2**). These differences cannot result from clonal diversification. Therefore, the diversity observed within the Silk group results from two different sexual events, probably from parents of similar genetic background.

#### Kalapua

The Kalapua group, characterized by an ABB genomic composition, is a popular cooking banana variety in Papua New Guinea. Within our sample set of five Kalapua accessions, we identified two distinct genomic mosaics. The first mosaic pattern was present in three of the samples, while the second pattern was found in the remaining two. The observed differences between these mosaics consist of varying proportions of A and B genomes in four specific genomic regions. The first telomere of chromosome 4 and the interstitial region of the first arm of chromosome 8 are A2:B1 in Kalapua-1 and A1:B2 in Kalapua-2. In addition, on chromosome 9, the first telomere is A1:B2 in Kalapua-1 and A0:B3 in Kalapua-2 while the second telomere is A2:B1 in Kalapua-2 and A1:B2 in Kalapua-1. Kalapua patterns may have resulted from the accumulation of mitotic homoeologous chromosome exchanges. However, mitotic recombination events are rare and these four cumulated events may alternatively have resulted from two sexual events among similar parents.

#### Pisang Awak

The Pisang Awak group comprised two triploid and three tetraploid patterns present in our sample. For the triploid patterns, the pattern Pisang-Awak-1 (PA-1), was found in 7 accessions while Pisang-Awak-2 (PA-2) was found in 2 accessions from India. Eight variations in the patterns of homologous exchanges between the A and B genomes were observed between these two mosaic patterns. Differences in A/B homoeologous exchanges consisted either in the presence or absence of these events or in variations in the size of common events. This finding is not consistent with two genotypes deriving from each other clonally and rather supports the idea that they were both sexually produced, probably from the same AB parent who produced recombined but unreduced (2x) gamete. Furthermore, A/B homoeologous exchanges enabled to hypothesize a pedigree relationship with the three tetraploids patterns identified in this cultivar group (4 samples). These four accessions, with ABBB genomic composition, were morphologically included in the Pisang Awak group and were produced from unreduced (3x) ABB triploid Pisang Awak gametes crossed with a haploid (1x) B gamete. The A/B introgressions patterns observed in the triploid and tetraploid Pisang Awak samples support pedigree relationships between PA-1 and ‘Ramu Yawa’ (Pisang-Awak-4x-1) from Papua New Guinea. Equally, direct ancestry can be inferred between PA-2 and ‘Pisang Awak’ (Pisang-Awak-4x-2) from Sri Lanka. The third tetraploid pattern, discovered in ‘Foulah 4’ and ‘Nzizi’ (Pisang-Awak-4x-3) from Ivory Coast and Nigeria respectively, did not correspond to any of the triploid Pisang Awak described, suggesting that at least a third triploid form may exist or may have existed (**Fig. 2**).

**Fig. 2.**
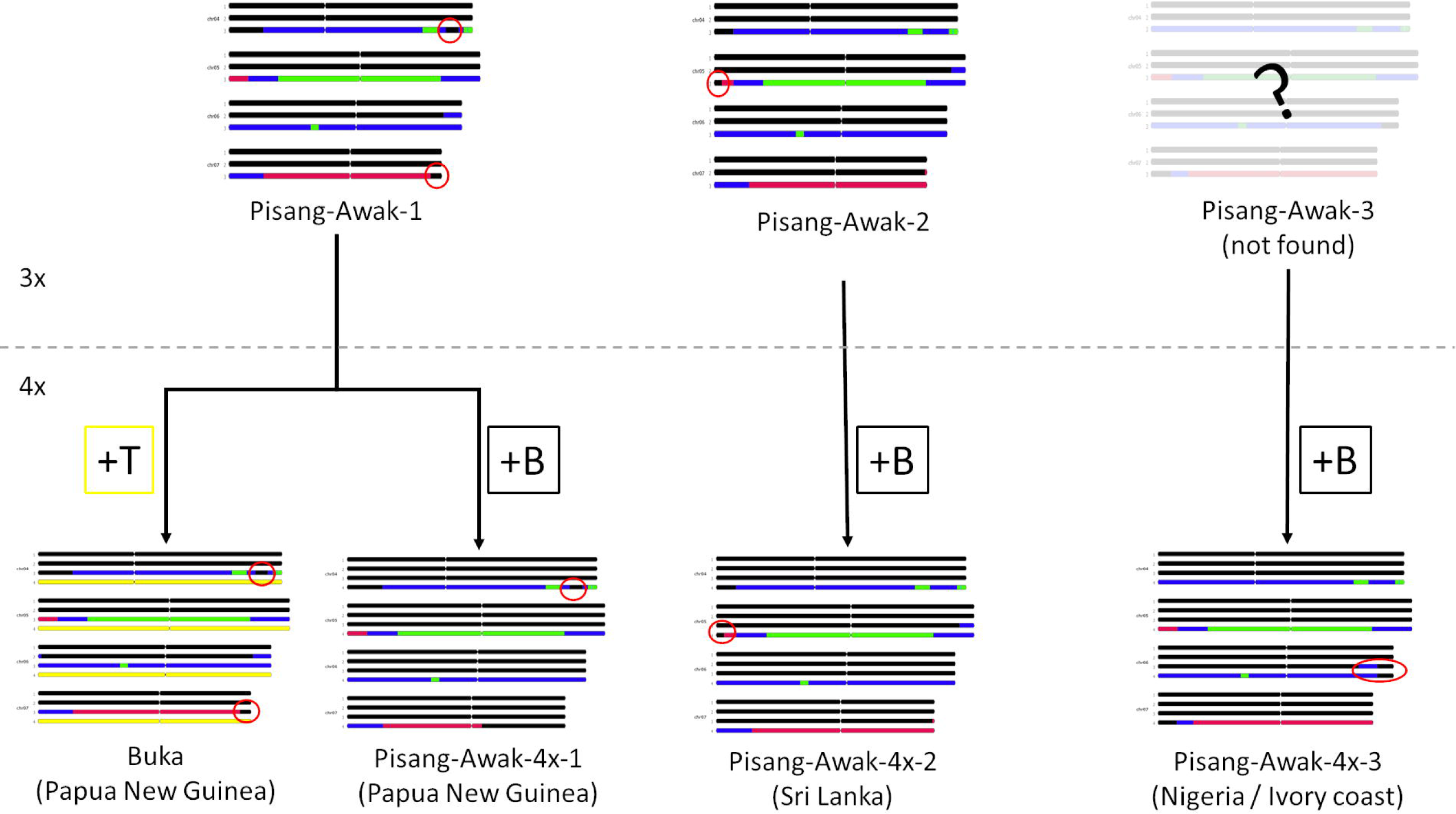
Genetic Diversification in the Pisang Awak Group. This illustration showcases the patterns of five Pisang Awak variants and a Pisang Awak-derived intersectional hybrid for four sets of chromosomes (chromosomes 4, 5, 6 and 7), each marked by distinctive events (red circle) inherited from one of the parents (other differences may result from recombination events that occurred during the production of unreduced gametes (3x). The integration of additional genomes into the triploid (3x) patterns (Pisang Awak-1/3), denoted by specific letters (B for balbisiana, represented in black; T for Australimusa, shown in yellow), has resulted in the formation of closely related tetraploid (4x) varieties. The triploid Pisang Awak-3 is represented with partial transparency as it was not found in the sample but could be inferred from its progeny. Coloured segments show the contribution of each of the ancestral genepools.

#### Kunnan and Ney Poovan

Two cultivar groups from India and with AB genomic composition are defined, the Ney Poovan and the Kunnan groups, but their morphological characteristics are not clear. In our sampling, 8 accessions with an AB genomic composition could be affiliated to either cultivar group. These accessions displayed four different mosaic patterns with at least one A introgression into their B genome (consistent with Cenci *et al*. 2021), and with a malaccensis ancestry dominance as A-donor genome. Two patterns were discovered in both Ney Poovan (3 accessions) and Kunnan (4 accessions) but the correspondence between the morphological assignation and the patterns was incomplete. Since these cultivar groups were not extensively documented and many synonyms and overlaps exist in India, the true assignation of the accession confusingly named ‘Kunnan’ (ITC1034) but classified as Ney Poovan, was difficult to assert.

### Other cultivars (clusters and individual accessions)

Some accessions of the set were ambiguous in classification, with morphological similarity with well-known cultivar groups, but not complying enough to discriminating criteria to be considered as part of these defined cultivar groups.

#### Similar to Mutika/Lujugira

Four accessions were collected in Tanzania with unclear classifications (De Langhe *et al*. 2001) and comprise two distinct mosaic patterns that share similarities with the Mutika/Lujugira group. Notably, they include a significant malaccensis component (12 centromeres and 6 centromeres, respectively), similar to what was observed in the Ilalyi group (**Table 2**). The first pattern, Kikundi, is identified in three accessions. ‘Ntebwa’ and ‘Ntindi I’, both from the Tanzania’s Usambara region differ in their uses, ‘Ntebwa’ is used for cooking, and ‘Ntindii I’ serves both as a cooking (flour) and dessert banana. The third accession ‘Kikundi’ differs by the pinkish colour of the pseudostem contrasting with the green observed in the two others. The second pattern, Luholole, is represented by a single accession from the Morogo district of Tanzania.

#### Similar to Plantain

Two accessions can be linked morphologically to the Plantain group. The accession ‘Kupulik’ was collected in the late 1980’s in Papua New Guinea (Island of New Ireland in the Bismark Archipelago) as a Horn type Plantain but it is not a Plantain. Its mosaic shares similarities with both Plantain and Iholena, but the malaccensis component is absent in ‘Kupulik’. Two other accessions originating from Papua New Guinea, ‘Bubun’ and ‘Navente 2’, exhibited the same pattern. Then, the cultivar **‘**Mnalouki’ from the Comoros (Perrier *et al*. 2019), shares two haplotypes with Plantain cultivars and was proposed to be a progeny of a Plantain (2x gamete) x Mchare (1x gamete) (Martin, Baurens, *et al*. 2023).

#### Similar to Iholena

Some level of morphological confusion exists around the Iholena group (Arnaud and Horry 1997; Kagy *et al*. 2016; Sachter-Smith *et al*. 2021; Sachter-Smith and Sardos 2021a). In our set, five accessions sharing some, but not all, morphological features of the Iholena exhibited two different and distinct mosaic patterns. The accessions ‘Rukumamb Tambey’, ‘Tigua’ and ‘Balabolo 1’ form the Rukumamb Tambey cluster. They share the bunch shape and the colour of the flesh with Iholena, but their fruits don’t turn yellow when ripe and the lower surface of their new leaf is green. The accessions ‘Arawa’ and ‘Mamae Upolu’ displayed a second mosaic pattern and were also different in their morphology (notably ‘Arawa’, which had an overall more diploid look at collect). ‘Mamae Upolu’, collected in Samoa, differs from Iholena by its slightly more upward fruits, the green lower surface of the new leaf and the more yellow colour of the flesh. Despite their morphological proximity, these two mosaic patterns differ from Iholena by the ancestry of 5 and 6 centromeres, respectively, and the notable presence of one malaccensis centromere that is absent in Iholena (**Table 2**).

#### Similar to Maia Maoli/P**ō**p**ō**ulu

We observed 6 bananas accessions that morphologically resemble the Maia Maoli/Popoulu (MMP) cultivars but with 3 different patterns. They were all collected in Papua New Guinea and surrounding islands and are composed of three different mosaic patterns that share a common background with the four MMP patterns previously identified and with other AAB cooking bananas. The first pattern, named here Wan Gevi, is composed of 2 accessions. The second pattern is represented by a unique accession ‘Buka Kiakiau’. Unlike the two other patterns, the third pattern named Ruango Block and made of 3 accessions, contains malaccensis as contributor (4 centromeres).

#### Clusters of accessions with distinctive morphotypes

We noted in our set some clusters of accessions, that may correspond to morphological variants of a same genomic pattern. It was notably the case of three sets of diploid accessions. The first set, composed of three accessions from the Philippines and Malaysia, was called here the Bata-Bata cluster. The second set was composed of four accessions from Papua New Guinea and was named here the Te’engi cluster. Then, the Talasea cluster was composed of two accessions collected in Papua New Guinea outer islands, one being likely the reddish variant of the other. Two morphological variants were also observed in the ABBT Buka cluster, ‘Buka’ being a shorter variety than ‘Bukayawa’.

### Individual profiles

Finally, we listed in the catalogue individual accessions which cannot be morphologically assigned to described cultivar groups and which have specific genomic patterns. Several diploids of AA genomic compositions presented interesting characteristics. The cultivar ‘Khai Na On’ is the male (1x gamete) parent of the ‘Gros Michel’ group (Raboin *et al*. 2005; Hippolyte *et al*. 2012; Martin, Cottin, *et al*. 2023). The cultivar ‘Pisang Madu’ is the keystone to the genome ancestry mosaic painting approach as it enabled the discovery and the identification of diagnostic SNPs for two uncharacterized ancestors of cultivated bananas and it is related to the Cavendish group. Its genome is indeed composed of a full haplotype of the unknown ancestor m1, proposed to be halabanensis from Indonesia (Martin, Cottin, *et al*. 2023). Additionally, it also contains 6 centromeres belonging to the unknown ancestor (Sardos *et al*. 2022; Martin, Cottin, *et al*. 2023). ‘Manang’, an accession from the Philippines, has a significant contribution from burmannica (4 centromeres), same as ‘Matti’ from India, also included in the catalogue with nearly a full haplotype of burmannica.

For the triploids, the ‘Chuoi Mit’ cultivar from Vietnam exhibits an ABB genomic composition, featuring an A genome that closely resembles the A genome found in the Bluggoe/Monthan pattern. The proximity of these cultivar groups has already been reported based on homoeologous exchanges between A and B subgenomes (Cenci *et al*. 2021) and is now confirmed analysing the A mosaic pattern. ‘Pisang Slendang’, an Indonesian AAB cultivar, shares one of its A haplotypes with both the Bluggoe and Monthan groups, as well as with ‘Chuoi Mit’. ‘Chuoi Xi Mon’ bears an unidentified genome differing from the characterized Unknown genepool.

The cultivar ‘Auko’ (synonym ‘Vunapope’) from Papua New Guinea, with an ABB genomic composition, has a unique A mosaic pattern among all cultivated bananas (no zebrina ancestry). It is only made of banksii and schizocarpa. This finding supports the hypothesis of the early domestication of banana in New Guinea from where *M.a.* ssp *banksii* and *M. schizocarpa* originated (Carreel *et al*. 2002) and as recently showed by chromosome painting (Martin, Cottin, *et al*. 2023). On the other side of the spectrum, the cultivar ‘La’ from Vietnam with an AAB genomic composition has 14 zebrina centromeres and only one small schizocarpa introgression. The ‘Ya Ta Na Thin kha’ accession collected in Myanmar was provided with a poor classification (*Musa*) and was identified as a triploid ABB in our analysis. Its genomic composition is rich in burmannica, like the Klue Teparod group. However, the pattern of ‘Ya Ta Na Thin kha’ is different, notably with a small introgression of zebrina in the first arm of chromosome 9 that is absent in Klue Teparod which exhibits a balbisiana introgression in the same region. These two patterns shared recombination breakpoints, indicating common evolutionary history, even suggesting that ‘Ya Ta Na Thin kha’ could be one of the genotypes at the origin of this cultivar group (**Fig. 3**). However, in the absence of morphological description available, we were not able to assess whether ‘Ya Ta Na Thin kha’ belongs to the Klue Teparod group.

**Fig. 3.**
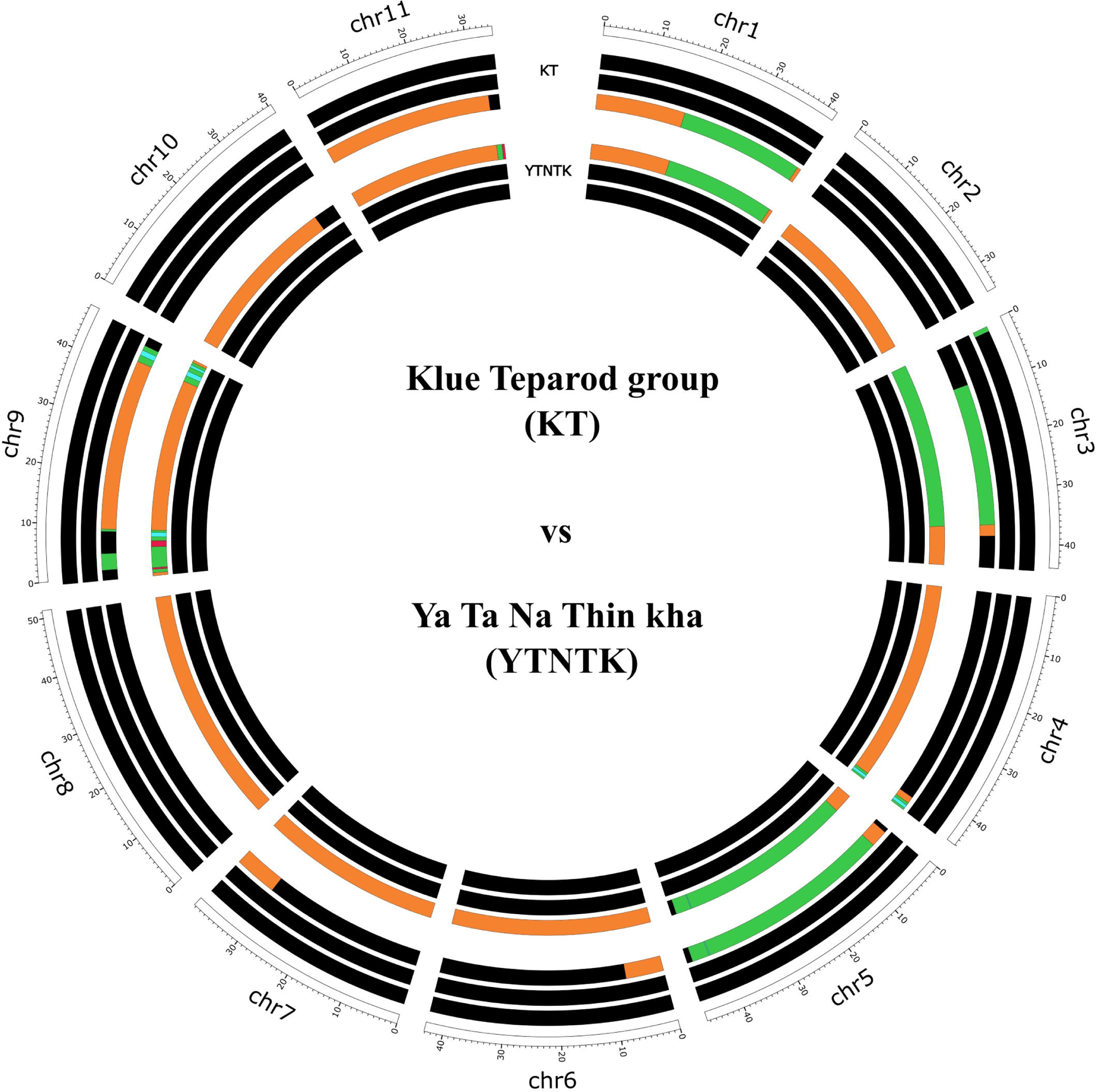
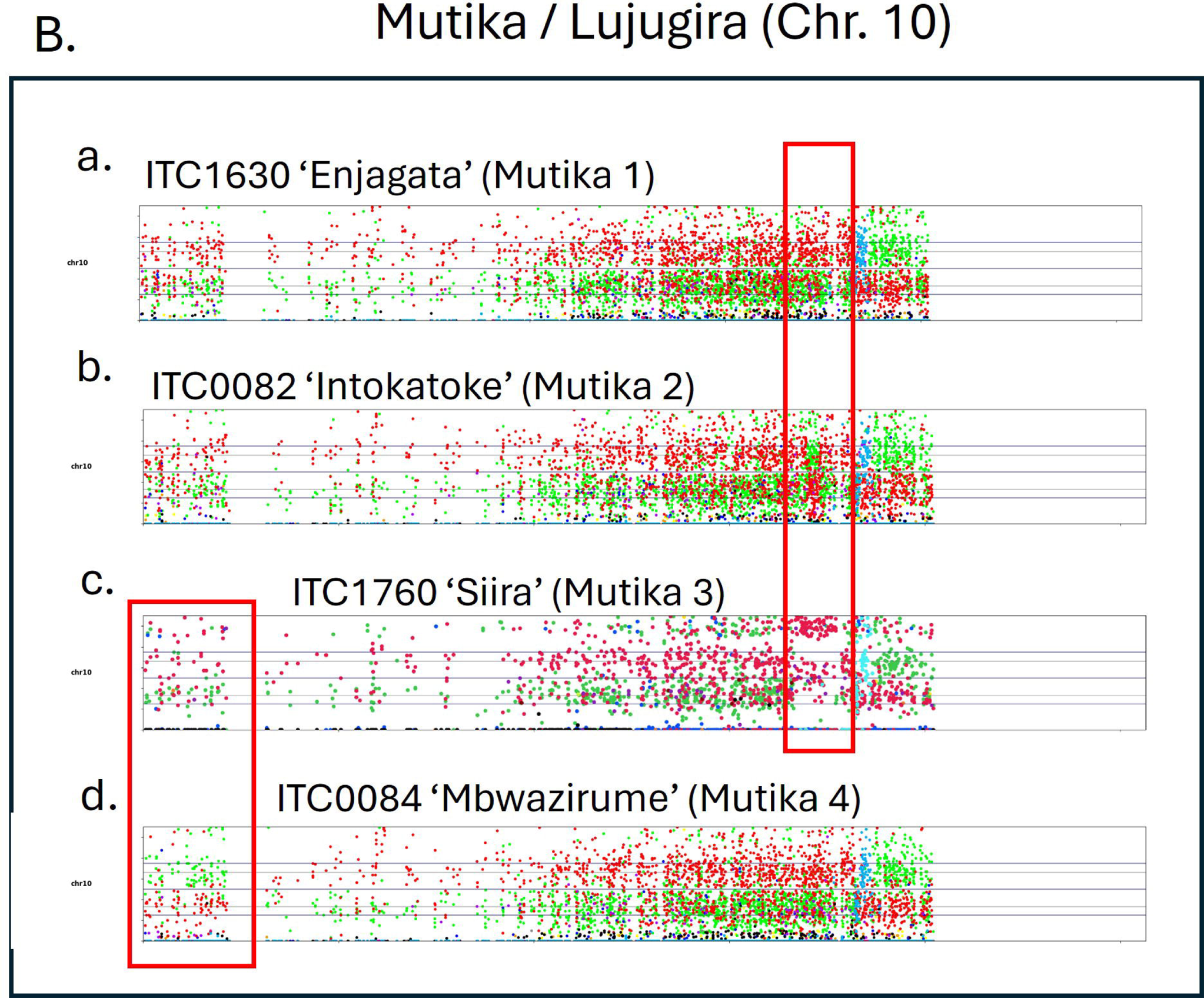
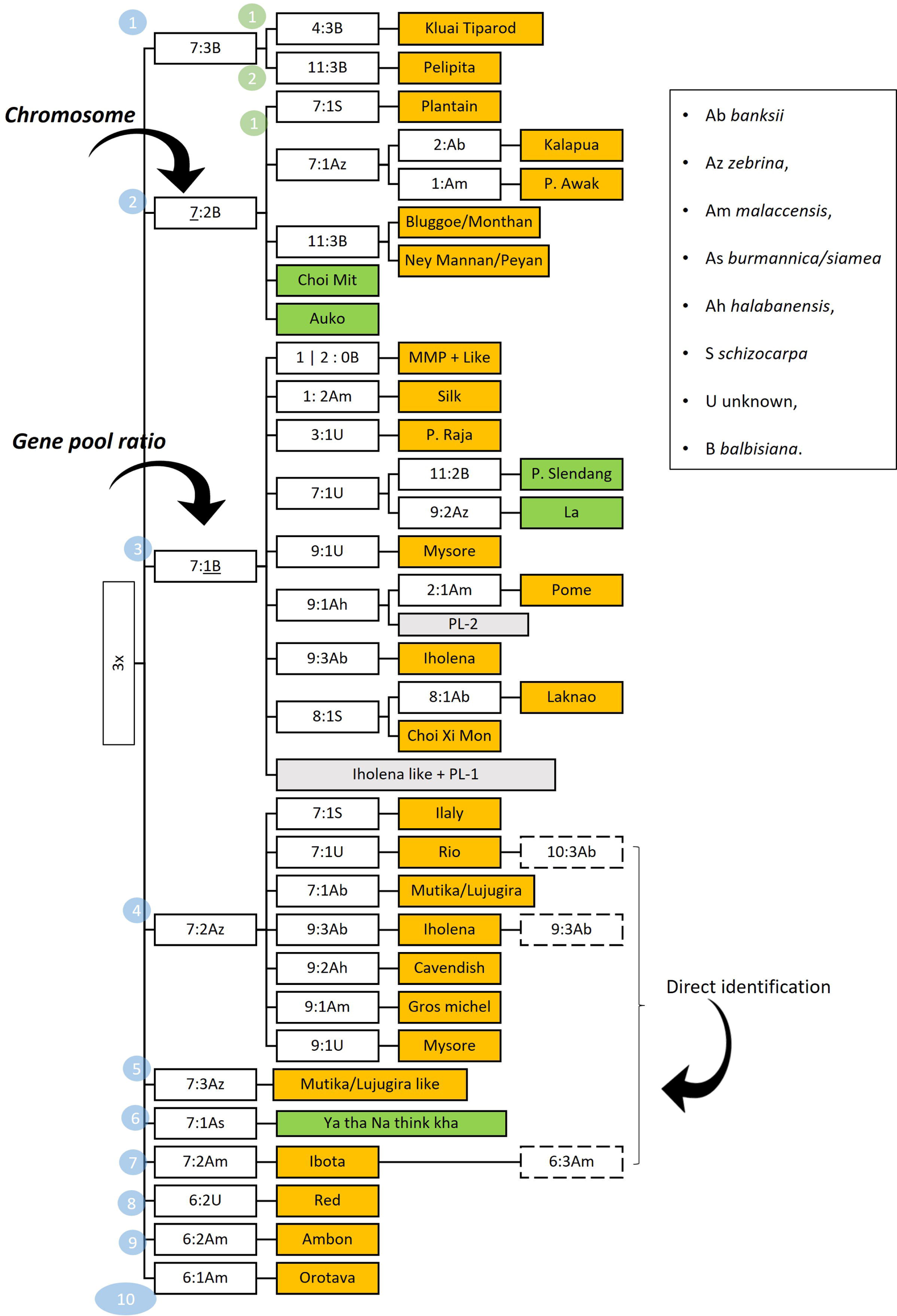
Genome ancestry mosaic painting applied to the Klue Tiparod (ABB, 3x) and ‘Ya Ta Na Thin kha’ (collected as Musa spp), revealing related pattern and a possible pedigree relationship. The colours of segments correspond to ancestral contributions (black: M. balbisiana, green: M. acuminata banksii genetic group, and orange: M. a. burmannica including ssp. siamea).

Until recently, only one triploid cultivar with a full S haplotype was known. The cultivar ‘Toitoi’ was collected in the island of Bougainville in Papua New Guinea. Its unrecombined schizocarpa haplotype suggested it resulted from an unreduced (2x) gamete with an AA genomic composition like many diploids AA cultivated in the country and a regular (1x) gamete S, probably from a wild specimen of *M. schizocarpa* endemic to New Guinea. A second cultivar with an AAS genomic composition named **‘**Waga’ was found since then, still in Papua New Guinea, and has a different mosaic pattern which interestingly shows two *M. acuminata* ssp. *banksii* introgressions on the chromosome 4 of the *M. schizocarpa* haplotype that probably results from a different type of cross (**Dataset S2**). Several tetraploid cultivars with a T genome were discovered in Papua New Guinea. However, the SNPs assigned to T are representative of the whole former Australimusa section of the *Musa* species, from which arose the Fehi bananas, with no indication on the specific species or genepool involved. Among these tetraploid cultivars, ‘Kalmagol’ which morphologically resembles the Silk group, had a genomic composition of AABT with the AAB genome that could correspond to the pattern Silk-2. Among the three patterns with an ABBT genomic composition, the cultivars ‘Buka’ and ‘Bukayawa’ displayed an ABB genome that could have derived from the Pisang-Awak-1 pattern (**Fig. 2**) while the cultivar ‘Bengani’ likely derived from the pattern Kalapua-2. In Cook Island, ‘Rekua’, a second ABBT cultivar with an ABB genome like the Kalapua patterns was discovered. It is different from ‘Bengani’, and its ABB genome may have arisen from Kalapua-1.

## Discussion

### Clonal diversification at the genomic scale

Domestication of banana was a gradual process in which the selection for edible pulp led to today’s parthenocarpic and highly sterile cultivars (Simmonds 1966). In this scenario, cultivar groups were expected to correspond to cultivars clonally derived from each other with the clonal accumulation of both point mutations and epigenetic variations as main mechanisms of diversification (Simmonds 1966). For example, despite the extremely high levels of intra-cultivar group phenotypic diversity observed in the Plantain and Mutika/Lujugira groups (Tézenas Du Montcel *et al*. 1983; Karamura *et al*. 1998), they were both found genetically homogenous (Noyer *et al*. 2005; Kitavi *et al*. 2016), but with significant levels of epigenetic variations (Noyer *et al*. 2005; Kitavi *et al*. 2020). Here, we identified larger genomic variations in these cultivar groups, fixed through vegetative propagation. Specifically, deletions were observed in the Mutika-3 and Plantain-2 patterns, and another event interpreted as a homologous exchange due to mitotic recombination, was inferred in the Mutika-2 and Mutika-4 pattern. The patterns Plantain-2 and Mutika-2 are in accordance with the findings of Martin, Cottin, *et al*. (2023) for these two cultivar groups but the high number of accessions analysed in our set enabled the discovery of other variants. In the Plantain group, the deletion, frequently observed in African accessions, is absent in the Asian accessions, suggesting that this event occurred after the introduction of the first Plantain cultivar(s) in Africa and supports previous hypotheses (Perrier *et al*. 2011; Langhe *et al*. 2015) on the origin of this cultivar group in South-East Asia. For the Mutika/Lujugira cultivars studied here, the two genomic variants characterized on chromosome 10 were found in only a small portion of the samples. However, most of the Mutika/Lujugira analysed were introduced to the ITC from Rwanda and Burundi and may not represent the entire diversity of this cultivar group. Further genomic characterization of the Mutika/Lujugira germplasm across a wider geographical range is recommended for future studies.

Within those two cultivar groups, we did not observe obvious correlations between the small genomic variations detected and the striking phenotypic features of these cultivar groups, a pattern also observed with epigenetic variations (Noyer *et al*. 2005; Kitavi *et al*. 2020). However, these events constitute valuable markers for tracing the evolutionary history of Plantain and Mutika/Lujugira. Additionally, these genomic variations notably generate gene copy number variations, as identified in multiple crops (Yakushiji *et al*. 2006; Stein *et al*. 2017; Gabur *et al*. 2019), and may be linked to interesting traits, such as diseases resistance, as identified in a few somaclonal variants of Cavendish exhibiting deletions on chromosome 5 (Hou *et al*. 2022). Therefore, the specific regions of chromosome 10, where structural variations were identified in the Mutika/Lujugira and Plantain groups, merits further investigation.

### Sexuality still matters in cultivated bananas

The diversity of genome patterns observed within several cultivar groups, including well-known cultivar groups such as Silk, Pome, Maia Maoli/Popoulu and Pisang Awak, seemed difficult to explain only by clonal diversification. The variations observed, with centromeres of different origins and/or the accumulation of high numbers of recombination, rather supports a meiotic origin to these differences. However, it is striking that the different patterns observed in these cultivar groups are so similar that they might be siblings from the same parental clones, as inferred for the triploid Pisang Awak accessions (**Fig. 2**). This important finding supports Kagy *et al*. (2016) who proposed an enlarged vision of cultivar groups and considered that sets of closely related cultivars could ensue from different sexual events within similar or closely related parents. In addition, the Pisang Awak example, with the occurrence of tetraploid siblings of triploid landraces, illustrates that original clones can also be sources of sexual diversification within cultivar groups (**Fig. 2**). Pisang Awak, are more prone to set seeds (Simmonds 1962) and the tetraploid cultivars observed here show that farmers continue to select new banana cultivars that arise accidentally from seeds, further contributing to expanding diversity.

Interestingly, several of the accessions not affiliated to any cultivar group brought interesting insights by showing the progressive incorporation of exotic genepools. Notably, in East Africa where the Ilayi group and two other patterns, Kikundi and Luholole, display a common genomic background with the Mutika/Lujugira group characterized by an important contribution of banksii and zebrina, but supplemented by malaccensis. Such pattern was also observed in ‘Mnalouki’, when compared to Plantain as well as in the Ruango Block cluster when compared to Maia Maoli/Popoulu and to some extent in Rukumamb Tambey and Arawa clusters when compared to Iholena. Considering that banana’s domestication centre was likely located in New Guinea island (Martin, Cottin, et al. 2023) where only *M. acuminata* ssp. *banksii* and *M. schizocarpa* can be found, the genomic constitution of ‘Auko’, free of zebrina genepool, suggests that the addition of zebrina to the genomic backgrounds of cultivars likely resulted from secondary diversification events. However, the common contribution of zebrina to all other cultivars suggests that the addition of zebrina precluded the insertion of malaccensis in banana cultivars. This scenario is consistent with the geographic distribution of *M. acuminata* subspecies and is in line with the correlation observed between the wild subspecies geographical ranges and the wild ancestors’ contribution to local cultivars (Sardos et al. 2022; Martin, Cottin, et al. 2023). Therefore, these accessions related to known cultivar groups but with additional contribution of malaccensis may result from more recent sexual diversification, such as ‘Mnalouki’ which may be a sibling of Plantain (Martin, Baurens, et al. 2023).

In addition, the tetraploid accessions with a T haplotype that were collected in the Pacific showed the incorporation of a supplementary genepool into cultivars from the Silk, Pisang Awak and Kalapua groups. Interestingly, the Silk and Pisang Awak groups originated in India and South-East Asia respectively, while wild and cultivated specimen of the former Australimusa section can be found only in an area going from Sulawesi (east Indonesia) to the Pacific Islands. It shows that these hybridizations occurred more recently, after the introduction of these cultivar groups in the distribution range of the ex-Australimusa specimens. It illustrates the importance of conserving local genetic resources as they can still be active in crop diversification.

### Implications for farming system, taxonomy, breeding and conservation

#### Farming systems

In clonal crops, vegetative propagation is an efficient way to preserve and multiply favourable genotypes that would not be maintained through sexual reproduction (McKey et al. 2010). For banana, farmers have historically selected and preserved varieties upon noticing changes in traits in the field, whether clonal (Karamura et al. 2010) or resulting from residual sexual events (De Langhe et al. 2010; Martin, Baurens, et al. 2023). Our hypothesis of genomic variations with two origins, clonal and sexual, co-existing in the overall diversity of cultivated bananas have implications in a context where monoclonal agriculture puts banana cultivation at risk in the face of biotic and abiotic stresses. These variations are valuable sources of diversity that can be overlooked by farmers. As stated before, deletions and duplications are sources of gene copy number variations that can result in differential phenotypes, just as homologous exchanges do. The use of this intra-cultivar group diversity would be an innovative way to diversify agrosystems by ensuring the co-existence of clonal and sexual variants in farmer cultivar portfolios. Since cultivar adoption is affected by a combination of sensory characteristics, agronomic properties and environmental and socio-cultural factors affecting production (Madalla 2021), planting different genomic patterns associated to the same cultivar group could allow to overcome part of these constraints while introducing diversity in farmers’ fields. Equally, the cultivars sharing morphological characteristics with known cultivar groups and which were found hybridized or introgressed with exotic genepools, could allow the introduction of additional genetic diversity, hence with putative beneficial new traits, while enhancing the likelihood of acceptance by farmers.

#### Classification of cultivated bananas

Our results show that intra-cultivar group diversification is made of a combination of sexual and clonal diversification and pleads for a relaxed definition of the cultivar group concept as the set of closely genetically related individuals sharing peculiar morphological characteristics. Classification criteria should be revised combining morphological assessment and chromosome painting results, for example using genomic determination keys (**Fig. S2**).

Our findings also raise new questions about the current taxonomy. For example, it may not be accurate to keep Bluggoe and Monthan as two separate cultivar groups while they share seemingly identical genomic backgrounds. Equally, the community should consider the creation of new cultivar groups as (clusters of) new genomic patterns are discovered. In some cases, cultivar groups may also be enlarged in a way they would incorporate the “grey zones” of cultivars resembling the "core” accessions of the cultivar groups but differing in their genomic patterns.

In such revision effort, some accessions would remain alone, constituting de facto cultivar groups with only one cultivar, this being relative to the current sample and possibly revisited with the addition of new cultivars. The catalogue presented here (**Dataset S1**), conceived as a dynamic and evolutive document, constitutes a valuable supporting tool for this task.

#### Breeding requires to maximize diversity

The chromosome painting approach was shown to be useful to understand the formation of current varieties (Martin, Baurens, *et al*. 2023), an essential point to ease successful breeding. It could also be a useful tool for breeders, both to support the selection of parents and the selection of hybrids (Cenci *et al*. 2023). The catalogue presented here, and the association of the patterns discovered in active genebank accessions, opens the door to an optimized use of banana diversity for breeding crosses. The occurrence of numerous individual accessions that cannot be affiliated to existing cultivar groups and that display unique and peculiar genomic background presents a fresh perspective for the use of original accessions as parents. Additionally, the intra-cultivar group diversity could also be a valuable resource for breeding. The selection of parents among the different clones or variants within a given cultivar group could allow the incorporation of new genetic variation in existing breeding schemes and may result in the incorporation of possible useful traits in the obtained progenies.

#### Let’s keep characterizing and collecting

Methods using chromosome characterization based on ancestral origin (mosaic genomes) were proven efficient to support germplasm characterization (Santos *et al*. 2019; Ahmed *et al*. 2019; Martin *et al*. 2020; Wu *et al*. 2021). The present study aimed at providing an efficient tool to support the community of banana researchers and workers in the task of classifying cultivars. This tool was also found helpful for the resolution of taxonomical issues that may arise. In this study, we found that about 25% of the taxonomic information displayed in the existing passport data required corrections, or clarifications when details were missing (**Table S1**). This catalogue also constitutes a tool to support the routine management of banana genebanks through molecular characterization. Combining high throughput genotyping with this tool constitutes a much faster way to classify germplasm when compared to morphological characterization. Additionally, creating a baseline by genotyping germplasm that enters collections can help track *in-vitro* induced aneuploids (found here but data not shown), synonyms and potential duplicates.

In this study we could not characterize the entire banana genepool. For example, the chromosome painting results obtained for the many unique diploid cultivars from Papua New Guinea are not presented here but may be subject to a separate study. Also, the characterization of the popular Saba cultivars from the Philippines was not conducted due to lack of material available in the genebank. Nevertheless, this catalogue constitutes a baseline that can be further enriched as prospections and genomic characterization continue. An online version aims to be dynamic, continually expanded, and updated as characterizations of the ancestral gene pools and sequencing technologies improve.

Importantly, we provided evidence for the richness of patterns identified in a small number of accessions and that much more diversity exists in regions that have not been explored or have been underexplored. Screening only a few new accessions from recent collecting missions was sufficient to discover new genetic profiles that did not match defined cultivar groups, maybe justifying the creation of new cultivar groups, or blurring the lines of the defined ones. Now that germplasm characterization has entered the area of genomics, it is more than likely that further prospection of banana diversity in farmers’ fields will enable the discovery of new variants and new genotypes. Obviously, gaps remain in our perception of banana diversity and additional collecting missions are necessary to enrich our understanding of banana diversity and to fill the gaps in the collections.

## Data availability

The sequencing reads were deposited in the NCBI SRA associated with the BioProject PRJNA450532. SNP datasets were recorded in a database browsable via a web application https://gigwa.cgiar.org/gigwa (Sempéré *et al*. 2019; Rouard *et al*. 2022).

## Acknowledgments

We thank the International Transit Center for providing banana samples, BGI and LGC for their services for the RAD sequencing, and the AGAP Institute’s genotyping platform for the GBS. We acknowledge the CRP-RTB and the Genebank Platform of the CGIAR for the funding of the genotyping experiments as well as the Crop Trust for their support to the collecting missions. This work was technically supported by the CIRAD—UMR AGAP HPC Data Centre of the South Green Bioinformatics platform. We are grateful to the taxonomic information provided by the Taxonomy Advisory Group of MusaNet (https://musanet.org/who-we-are/taxonomic-advisory-group) for providing on regular basis critical insight to verify the true-to-typeness of the accessions.

We dedicate this work to two eminent banana scientists who passed away during the preparation of this manuscript. Professor Edmond De Langhe devoted his life and passion to bananas until the end and leaves an unvaluable contribution to our understanding of this crop. Dr. Hughes Tezenas du Montcel, a renowned expert in banana research, significantly enriched the field through his extensive collection of banana plants and laid the groundwork for the Musa Germplasm Information System. This work could not have been possible without their dedication to banana germplasm collection and conservation.

## Authors contributions

Conceptualisation, JS, AC and MRo; Provided material, IVdH, JP, WW; DTH; Library preparation and GBS genotyping: RR; Passport Data Curation, JS, CJ, RC, MRu, YM, GSS, MRo; Methodology and bioinformatic Analysis, GM, CB; Mosaic analyses and curation, AC, JS; Catalogue design, MRo, JS, Online catalogue development, VG; Data release in MGIS/Gigwa, CB, MRo; Writing – Original Draft Preparation, JS, MRo; Writing – Review & Editing, XP, GM, NY, CJ, RC, GSS, AC, NR; Funding Acquisition, AD, NR; All authors have read and agreed to the published version of the manuscript.

## Supplementary material

**Dataset S1**. Catalogue of cultivated bananas

**Dataset S2.** Additional mosaics for individual accessions

**Table S1.** List of plant materials used with consensus passport data from MGIS, morphological and genomic characterization.

**Fig. S1.** Examples of genomic events linked to clonal diversification inferred from chromosome painting raw outputs of VCFHunter. Colours correspond to *M. acuminata* ssp. *banksii* (green), ssp. *zebrina* (red) and *M. schizocarpa* (light blue). Plots of diagnostic SNPs on chromosome 10 of the Mutika/Lujugira group show a. a regular and most common profile, b. a switch in allelic ratio between zebrina and banksii in the interstitial region of second arm, c. a deletion of a fragment of the banksii haplotype in the interstitial region of second arm, and d. a switch in allelic ration between zebrina and banksii on the first telomere.

**Fig. S2.** Tentative genomic determination keys for accessions used in the molecular catalogue. In white rectangles are indicated Ancestral contributor and its ratio at the centromeric region (e.g. 7:3B for 3 balbisiana centromeric regions on chromosome 7), which are used to discriminate between cultivar groups and individual patterns (green rectangles). The diagram must be read left to right, top to bottom, as numbered.

