## Supplementary material for "Painting the diversity of a world’s favourite fruit: A next generation catalogue of cultivated bananas": Data S1

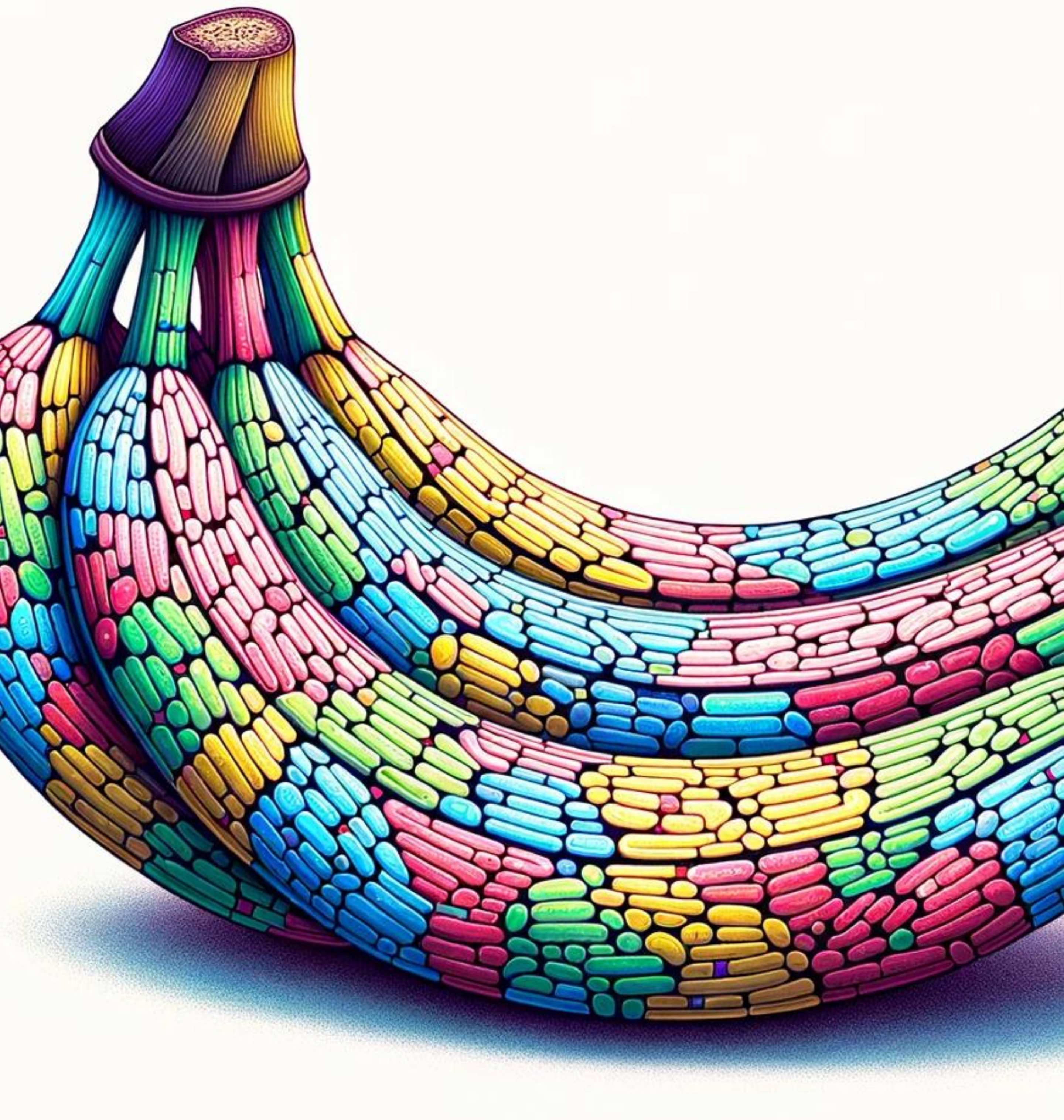

### A Genomic Catalog of Cultivated Bananas

Alliance Bioversity & CIAT

MusaNet

cirad

SAMPLE ONLY

**Coordination catalogue and editing:** Julie Sardos and Mathieu Rouard

**Passport data:** Julie Sardos, Mathieu Rouard, Christophe Jenny, Max Ruas with support of the Taxonomy Advisory Group (TAG)

**Photos:** Yaleidis Mendez, Christophe Jenny, Gabe Sachter-smith, Julie Sardos, Jorge Vargas, Miguel Dita, Lucien Ibonbondji, Lorna E. Herradura,

**Karyotypes:** Alberto Cenci, Julie Sardos, Guillaume Martin, Catherine Breton and Mathieu Rouard

**Graphic design and covers:** Mathieu Rouard and DALLE

**Citation:**

*Julie Sardos, Alberto Cenci et al. Painting the diversity of a world's favourite fruit: A next generation catalogue of cultivated bananas (2024)*

### FOREWORD

**Introduction**

Banana is an important food crop cultivated in many tropical and subtropical regions around the world. With the recent advances in genomics, a new powerful tool was developed enabling the fine-scale characterization of banana's ancestry along chromosomes, i.e. chromosome painting.

This catalogue, although not exhaustive, aims at providing an efficient tool to support the banana research and worker community in the classification of cultivars. It can be used as a tool to support the management of banana collections, which is a much faster way to classify them than with morphological characterization.

**Methodology**

We applied this method to a high-throughput genotyping data set obtained from 317 banana accessions spanning most of the known cultivar groups. This set included both genebank and new uncharacterized materials. By comparing curated morphological assignation to the chromosome painting results, we were able to compile a catalogue referencing the chromosome painting patterns of most of the described cultivar groups. In this approach, chromosome segments are colored according to their inferred ancestral genome contribution. The colors of segments correspond to the following ancestral contributors: black: M. balbisiana (B genome), pale-blue: M. schizocarpa (S genome), green: M. acuminata ssp banksii / *M. acuminata* ssp errans (A genome); blue: M. a. malaccensis (A genome) red: M. a. ssp zebrina (A genome), pink: uncharacterized genepools, and purple: M. a. ssp halabanensis, Orange: Musa a. ssp burmannica / Musa a. ssp siamea, Yellow: Austalimusa (T genome) (new Callimusa section ).

**Disclaimer**: The genome ancestry mosaics method proved to be extremely useful but is not exempted of methodological bias. The size of segments is approximate since it was calculated with high density markers and their arrangement established on parsimony and may not reflect the real haplotypes.

### RESOURCES USED


1. Daniells J. 2001. Musalogue: a catalogue of Musa germplasm: diversity in the genus Musa. Bioversity International.

2. Kepler, Angela Kay, and Francis G. Rust. 2011. World of Bananas in Hawai'i: Then and Now : Traditional Pacific & Global Varieties, Cultures, Ornamentals, Health & Recipes. Pali-o-waipi'o Press.

3. Irish, B.M., C. Rios, N. Roux, J. Sardos and R. Goenaga. 2016. Characterization of the Musa spp. Taxonomic Reference Collection at the USDA-ARS Tropical Agriculture Research Station

4. Martin G, Cardi C, Sarah G, et al. 2020. Genome ancestry mosaics reveal multiple and cryptic contributors to cultivated banana. The Plant Journal 102: 1008–1025.

5. Martin G, Cottin A, Baurens F-C, et al. 2023. Interspecific introgression patterns reveal the origins of worldwide cultivated bananas in New Guinea. The Plant Journal 113: 802–818.

6. Martin G, Baurens F-C, Labadie K, et al. 2023. Shared pedigree relationships and transmission of unreduced gametes in cultivated banana. Annals of Botany 131: 1149–1161

7. Musapedia, the banana knowledge compedium, https://www.promusa.org/Musapedia

8. Ruas M, Guignon V, Sempere G, et al. 2017. MGIS: managing banana (Musa spp.) genetic resources information and high-throughput genotyping data. Database: The Journal of Biological Databases and Curation 2017: bax046. https://www.crop-diversity.org/mgis/

9. Sachter-Smith, G.; Sardos, J. (2021) Bananas of Cook Islands: A catalog of banana diversity seen on the islands of Rarotonga and Aitutaki. Rome (Italy) : Alliance of Bioversity International and CIAT, 40 p.

10. Sachter-Smith, G.; Paofa, J.; Rauka, G.; Sardos, J.; Janssens, S. (2017) Bananas of the Autonomous Region of Bougainville: a catalog of banana diversity seen on the islands of Bougainville and Buka, Papua New Guinea. 97 p.

11. Sardos, J.; Paofa, J.; Janssens, S.; Sachter-Smith, G.; Rauka, G.; Roux, N. (2017) Banana collecting mission in the Autonomous Region of Bougainville (AROB), Papua New Guinea. 26 p.

12. Sreejith, P. E. and M. Sabu Edible Bananas of South India. Taxonomy and Phytochemistry. 2017. illus.(col.). 292 p.

13. Valmayor, RV, B Silayoi, SH Jamaluddin, S Kusomo, RRC Espino, and OC Pascua. 1991. Banana Classification and Commercial Cultivars in Southeast Asia.

14. Valmayor, R. V., R. Espino, and O. C. Pascua. 2002. "Wild and Cultivated Bananas of the Philippines.

### CULTIVAR GROUPS


*AA genomic composition*

**Mchare**........................ 8

**Pisang Jari Buaya**........ 9

**Pisang Lilin** ................ 10

**Sucrier** ........................ 11

*AAA genomic composition*

**Cavendish** .................. 12

**Gros Michel** ............... 13

**Red** ............................ 14

**Mutika/Lujugira** ....... 15

**Ilaly** ........................... 16

**Ambo** ........................ 17

**Orotava** ..................... 18

**Rio** ............................ 19

**Ibota** ......................... 20

*AB genomic composition*

**Kunnan** ...................... 21

**Ney poovan** ............. 22

*AAB genomic composition*

**Plantain** ....................... 23

**Maia Maoli-Popoulu** .. 24

**Iholena** ........................ 25

**Laknau** ......................... 26

**Pome** ............................ 27

**Mysore** ......................... 28

**Silk** ............................... 29

**Pisang Raja** .................. 30

ABB genomic composition

**Pisang Awak** ............... 31

**Bluggoe** ...................... 33

**Monthan** ..................... 34

**Ney Mannan** .............. 35

**Peyan** .......................... 36

**Kalapua** ...................... 37

**Klue Tiparot** ............... 38

**Pelipita** ....................... 39


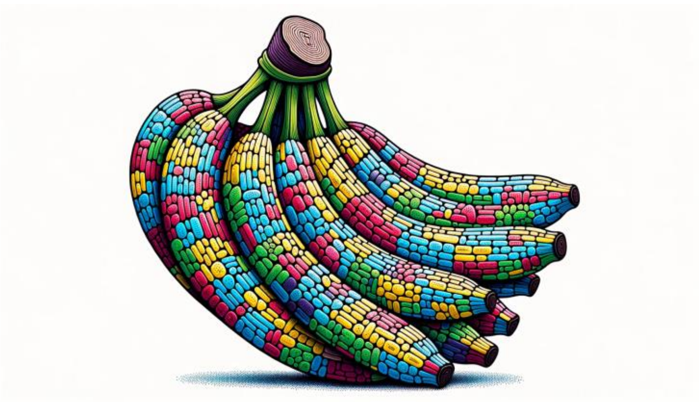

### CLUSTERS OF ACCESSIONS

*Similar to Mutika/Lujugira*
**Ntebwa** ............................. 41
**Luhulole** .......................... 42

*Similar to Plantain*
**Kupulik** .............................. 43
**Mnalouki** .......................... 44

*Similar to Maoli-Popoulu*
**Wan Gevi** ......................... 45
**Buka Kiakiau** .................. 46
**Ruango Block** ................. 47

*Similar to Iholena*
**Rukumamb Tambey** .......... 48
**Arawa** ................................. 49

*AA genomic composition*
**Bata Bata** ........................... 51
**Talasea** ............................... 52
**Te'engi** ................................ 53

*ABBT genomic composition*
**Buka** ................................... 54

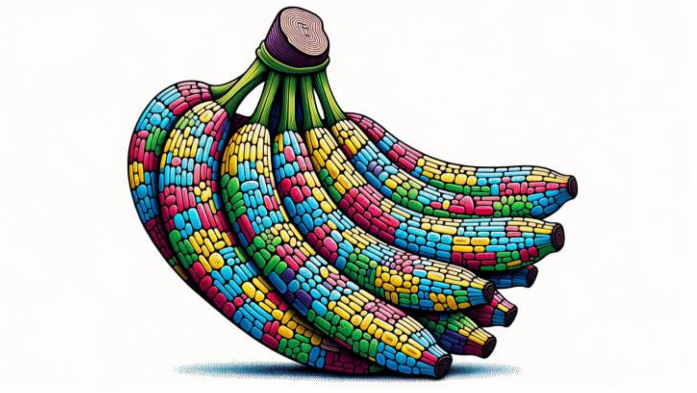

### INDIVIDUAL ACCESSIONS

*AA genomic composition*
**Khai Na On** ..................... 56
**Manang** .......................... 57
**Matti** .............................. 58
**Pisang Madu** .................. 59
**Pisang Pipit** .................... 60

AAS genomic composition
**Toitoi** ............................. 61

AAB genomic composition
**La** ................................... 62
**Pisang Slendang** ........... 63

ABB genomic composition
**Auko** ................................ 64
**Choi Mit** .......................... 65
**Ya Ta Na Thin kha**........... 66

ABX genomic composition
**Chuoi Xi Mon** ................. 67

*AABT genomic composition*
**Kalmagol** ........................ 68

*ABBT genomic composition*
**Bengani** .......................... 69
**Rekua** ............................. 70

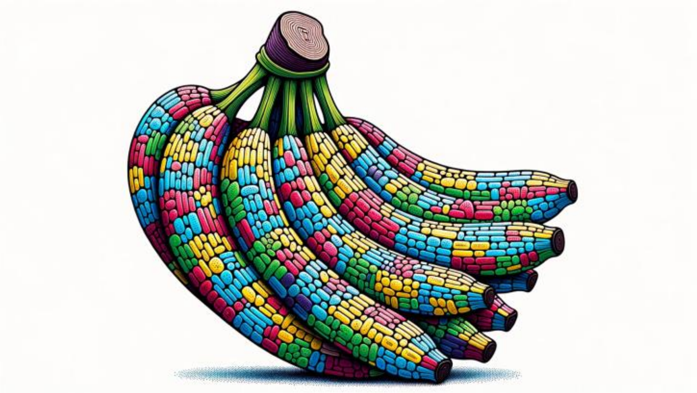

### Cultivar groups

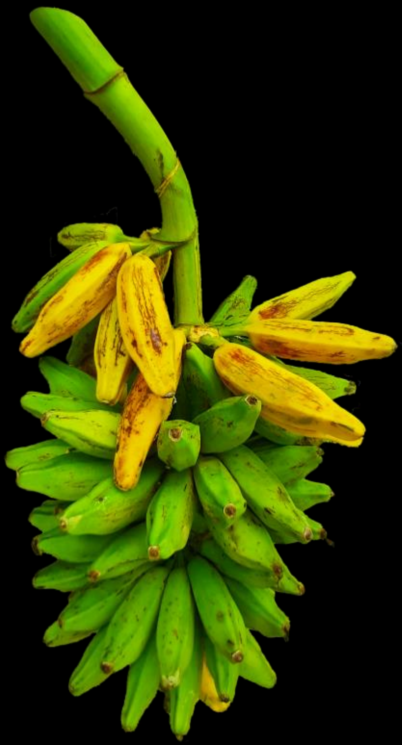

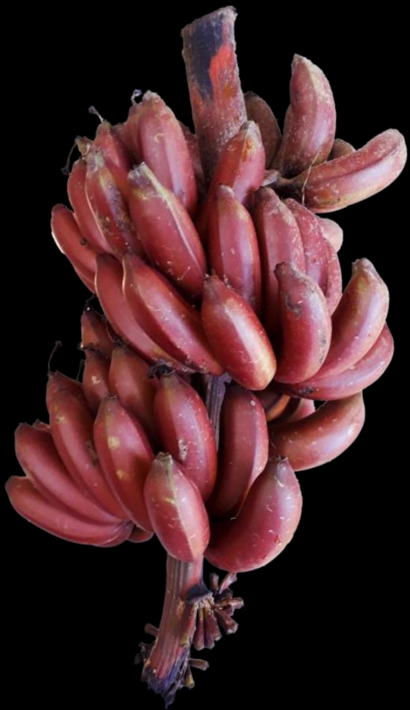

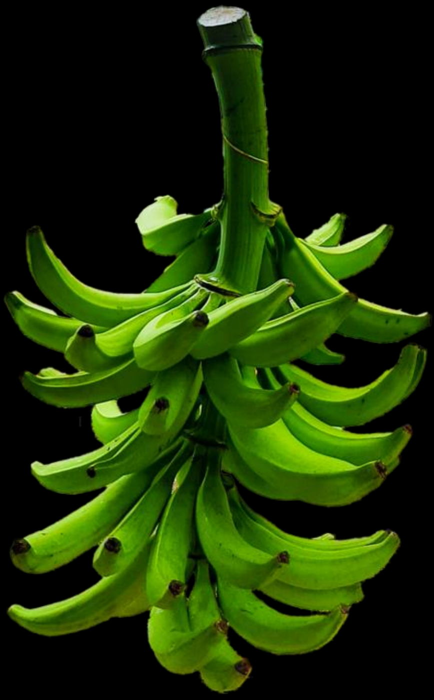

### Red group

#### Passport Data

**Classification**: Red (AAA)

**Biological status**: Cultivated

**Ploidy**: Triploid (*3x* = 33)

**Main distribution area**: Tropics and subtropics

**Uses**: Dessert

**Notes**:

- Most of the plant is red
- A green variant of Red exists, it is named Green Red
- Synonym : Figue rose

**Genomic features**:

- Two schizocarpa centromeres on chromosome 2, like Ibota and Maia Maoli Popoulou
- High contribution of unknown genepool

#### Morphological Characterization Pictures

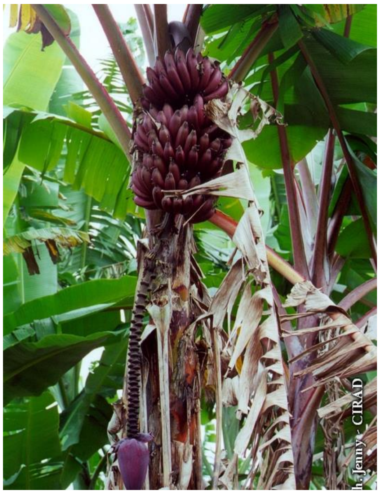

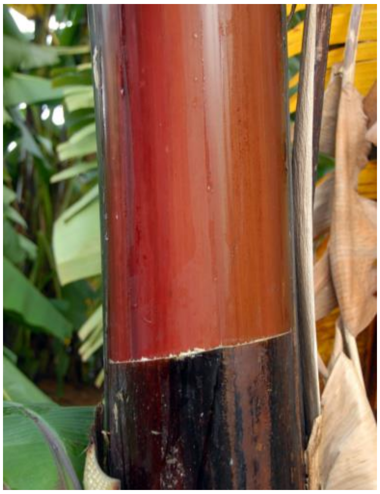

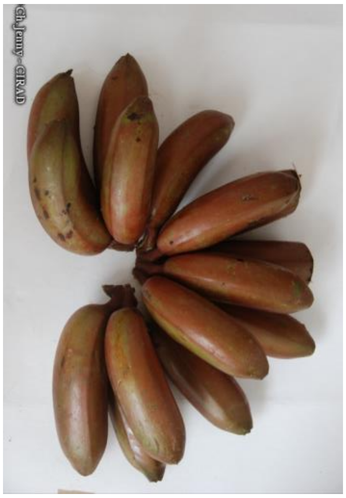

#### Molecular Characterization

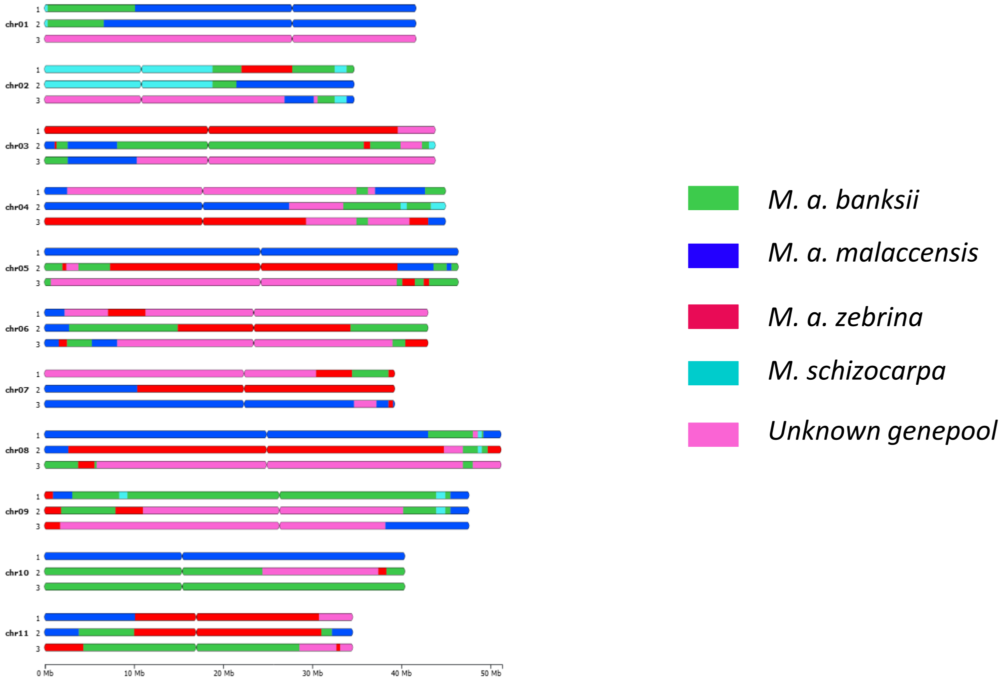


ITC1833 Shwe Ni

### Pisang Jari Buaya group

#### Passport Data

**Classification**: Pisang Jari Buaya (AA)

**Biological status**: Cultivated

**Ploidy**: Diploid (2x = 22)

**Main distribution area**: Asia and Pacific

**Uses**: Dessert, Breeding

**Notes**:

- Shape of bananas reminding crocodile's fingers. Rachis is full of neutral flowers with persistent bracts above the male bud. No pollen and seedless bunches
- Resistance to burrowing nematodes.

**Genomic features**:

- One haplotype from *M. a. halabanensis*

#### Morphological Characterization Pictures

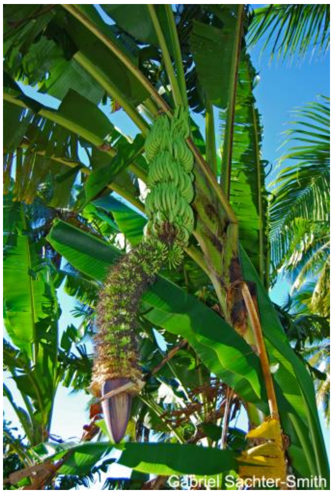


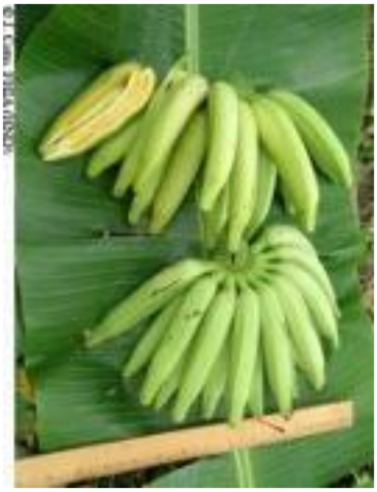

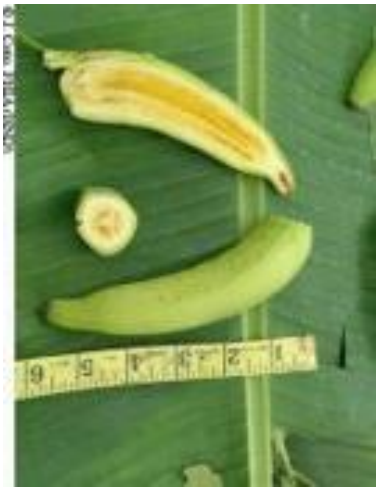

#### Molecular Characterization

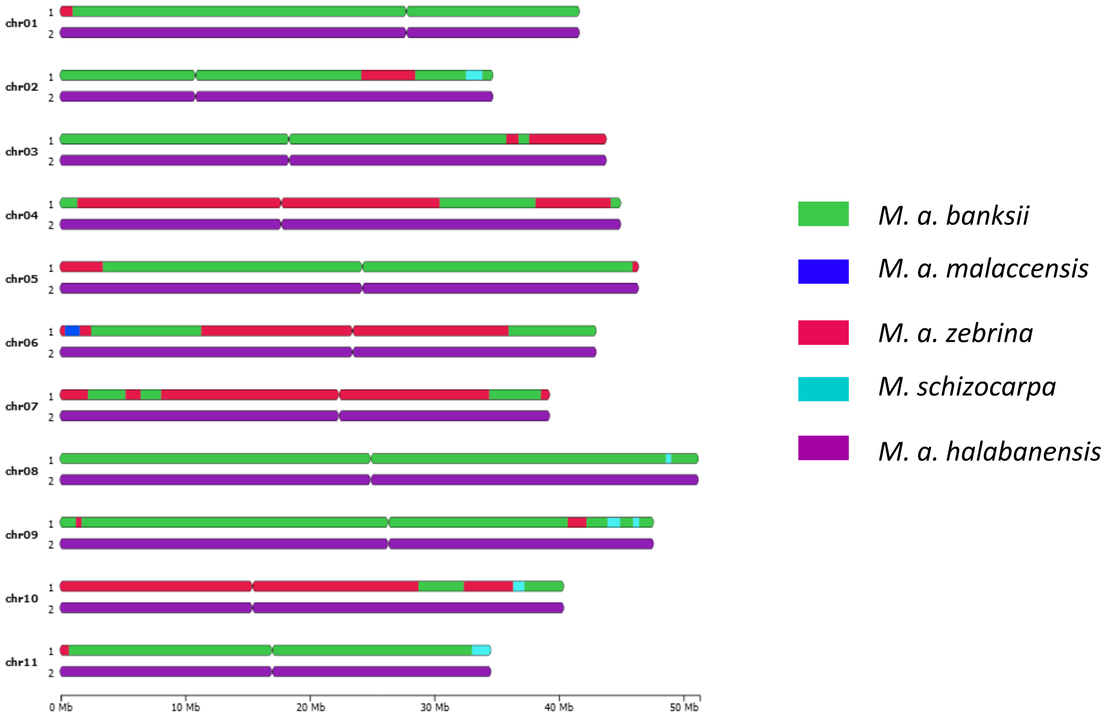


ITC0312 Pisang Jari Buaya

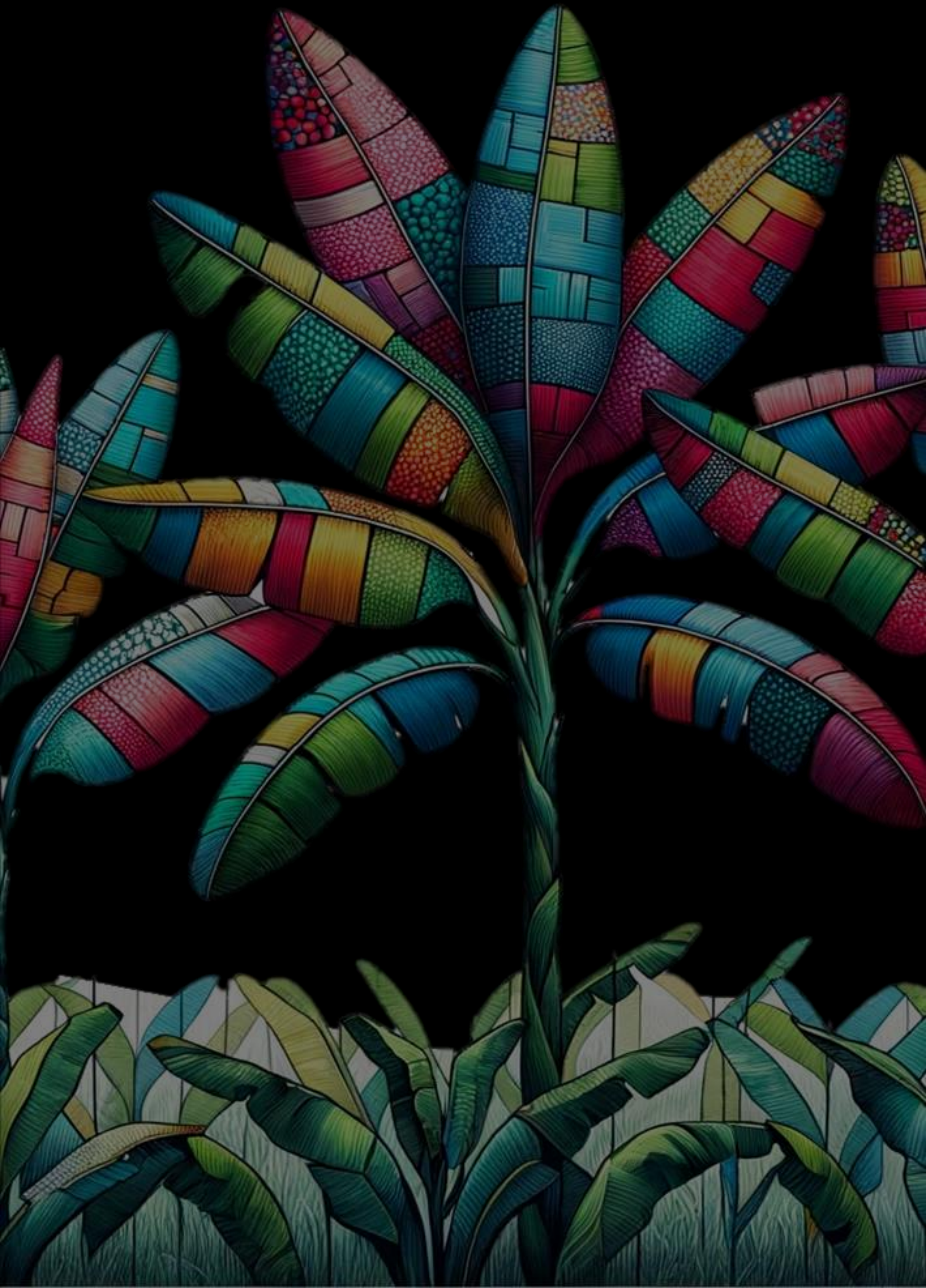
